# Characterizing Highly Conserved Fragments in 3’UTRs via Computational and Transfer Learning Approaches

**DOI:** 10.64898/2026.01.19.700376

**Authors:** Eric S. Ho, Ash Baeck-Hubloux, Nathan Dinh, Ava Severino, Ciara Troy

**Affiliations:** Department of Biology, Lafayette College, Easton, Pennsylvania, 18042, USA; Department of Computer Science, Lafayette College, Easton, Pennsylvania, 18042, USA

## Abstract

3’ untranslated regions (3’ UTRs) serve as regulatory platforms that modulate translation, mRNA localization, and stability through the binding of regulators, such as RNA-binding proteins (RBPs) and miRNAs, in a sequence-specific manner. These vital binding sites are often identified through orthologous regions among species. A separate but related discovery is the ultraconserved elements (UCEs) detected in human, rat, and mouse genomes two decades ago. However, our knowledge about their functions is limited. Perplexingly, alterations in UCEs in mouse embryos can still produce viable progeny with no observable phenotypic differences. The majority of UCEs are non-coding, though ∼8% are located in the 3’UTRs. Given the importance of 3’UTRs in gene regulation, we use a computational approach to identify highly conserved fragments (CFs) in 3’UTRs across diverse mammals, applying criteria appropriate for 3’UTRs (250 bp and ≥90% identity). Results show that they are not composed of simple repeats or low-complexity regions common to mammalian genomes. Using a transformer-based foundational genomic model, CFs are characterized as A and T-rich and distinguishable from the 3’UTR background. 36 human CFs from 25 genes are significantly depleted in variations in humans. They are enriched in neuronal tissues and play roles in neurodevelopment and RNA processing, mediated by RBPs and miRNAs. Our findings expand on existing studies that attribute UCEs primarily to enhancer function, suggesting a new path to explore the biological roles of UCEs in 3’UTRs.

**GRAPHICAL ABSTRACT:** 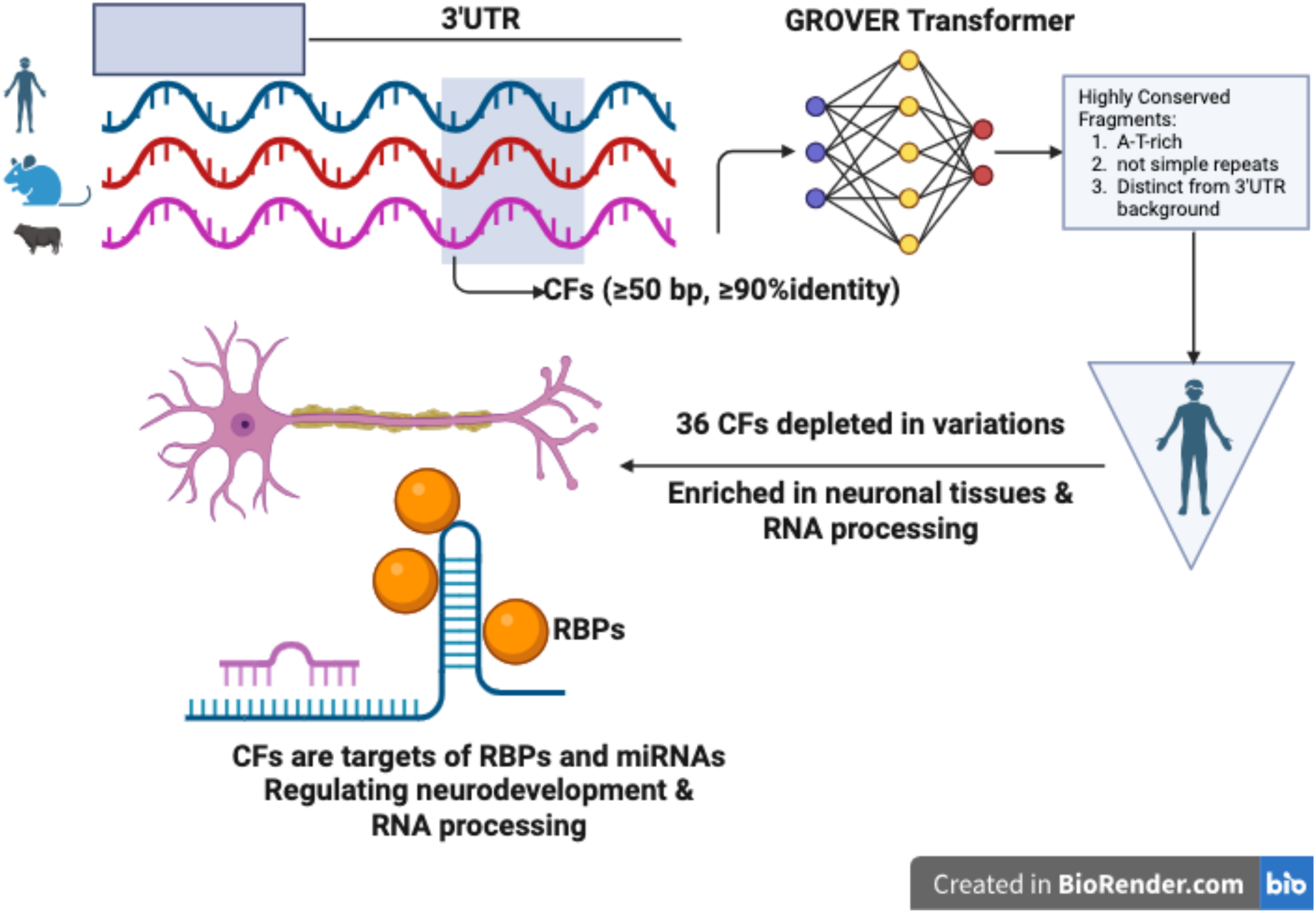

Created in BioRender. Ho, E. (2026) https://BioRender.com/dcyrx5f

## INTRODUCTION

3’ untranslated regions (3’ UTRs) harbor diverse regulatory elements that modulate essential biological functions, including mRNA stability, mRNA localization, and translation (reviewed in (1)). These functions are delivered through sequence-specific binding of RNA-binding proteins (RBPs) to these regulatory elements, followed by the recruitment of effectors by the bound RBPs. Notably, different combinations of RBPs and effectors can produce diametrically opposite effects. For example, AU-rich binding proteins TTP and KHSRP can mediate exosome complex binding, leading to mRNA degradation (2). On the other hand, the AU-rich-binding protein HuR (or ELAVL1) is unable to bind the exosome complex, thereby shielding HuR-bound mRNA from degradation (3). Post-transcriptionally, 3’UTRs contain sequence elements targeted by miRNAs (4, 5), thereby attenuating translation and mRNA stability (6, 7).

Moreover, alternative polyadenylation is prevalent in human and mouse genes, ∼54% and 32%, respectively (8), creating an additional mechanism to regulate genes through the shuffling in and out of regulatory elements in the long and short 3’ UTR isoforms, even if the coding sequence remains intact (9). For example, CD47 localization is regulated by alternative polyadenylation. The CD47 proteins expressed from long 3’ UTRs (∼3.5 kb) are anchored on the cell membrane, while CD47 proteins from the short 3’ UTR isoform (∼300 b) are localized mainly in the endoplasmic reticulum (10).

It has been established that non-coding variants are associated with diseases (11–13). There were high-throughput assays developed to screen for variants in 3’ UTRs (14). A cancer mutation burden metric that solely focuses on untranslated variants (15). These efforts underscore the critical regulatory roles of 3’ UTRs, implying that investigating the conservation of their sequences may offer insights into their functions. Unlike protein-coding regions (CDSs), 3’ UTRs are largely unconserved. Regulatory RNA elements in the 3’ UTRs are usually short (<10 bp). For instance, the motif of class I AU-rich element is AUUUA, and class II is UUAUUUA(U/A)(U/A). That said, we have discovered long (>50 bp) conserved fragments (>90% identity) in the 3’ UTRs among diverse mammals (human, mouse, rat, cow, and platypus), leading to the question of whether their conservation underpins crucial fundamental functions prevailing in mammals, or even vertebrates.

Previous studies have reported the presence of DNA ultraconserved elements (UCEs) in mammals and vertebrates (16, 17). Close to 500 UCEs were first reported in human, mouse, and rat, all of which are identical in length > 200 bp. With the availability of more genomes, subsequent studies have found over 14, 000 UCEs > 100 bp in at least three of the five mammals (human, mouse, rat, cow, and dog). However, only ∼8% of UCEs are reported in the 3’ UTR (18).

UCEs are found predominantly in non-coding regions, including introns, splice sites of protein-coding genes, and lncRNA exons (reviewed in (19)). Some UCEs are even found in the intergenic areas, though it is unclear whether they are transcribed (20). Their peculiar conservation has prompted speculation about their vital functions. However, the current picture remains perplexing. Enhancer is one of the functions being studied experimentally (21, 22), although perfect sequence identity is not required for transcription factor binding. A study found that alternation of ultraconserved enhancers produced no overt phenotypes (23). A recent study has reported that somatic mutations of certain non-coding UCEs are common in cancers (24). Genes possessing UCEs are frequently found to play roles in skeletal muscle and neurological development (22, 25).

Given the essential role of 3’ UTRs in gene regulation and the puzzling view of UCEs, both the author (26) and others (27) have reported the presence of highly conserved elements in the 3’UTRs in vertebrates. With the advances of deep learning, we aim to offer a renewed characterization of the conserved fragments (CFs) present in 3’ UTRs. By taking a contrastive approach, this study has revealed fundamental differences between CFs and their flanking unconserved 3’ UTR sequences. Such understanding might reveal obscure functional elements for RBPs and miRNAs. Although the CFs reported here show >90% identity, rather than 100% as reported in previous UCE studies, we argue that this threshold aligns with the nature of 3’UTR regulatory elements in which sequence variability is permissible for factor binding, such as RBPs and miRNAs. Such a view is also supported by previous studies (25, 28–30). To ensure the CFs reported here are not spurious, we implemented stringent filtering criteria (detailed below).

With the advanced foundation models in genomics, we used the GROVER model (31) to build a binary classifier to distinguish CFs from 3’ UTR sequences harboring no CFs, namely non-CFs. Importantly, by interpreting the model’s internals, i.e., the feature weights, it uncovered subtle sequence properties pertaining to CFs, shedding light on the unique propensities of CFs. In this article, we examine CFs from three perspectives: first, nucleotide propensities, complexity, and distinctive sequence elements. And then we focus on human CFs that are depleted of variations. Lastly, we explore the shared tissues, processes, and functions of genes harboring such CFs. Results suggest that the nucleotide composition of CFs differs from that of 3’UTRs, that genes harboring CFs are enriched in neuronal tissues, and that their functions are associated with neurodevelopment and transcription. Our findings guide further investigation and characterize functional roles of CFs in 3’UTRs.

## MATERIAL AND METHODS

### Untranslated Regions and Coding Sequences

We selected five diverse mammalian species to form the core group for analysis: Homo sapiens (human, 9606), Mus musculus (mouse, 10090), Rattus norvegicus (rat, 10116), Bos taurus (cow, 9913), and Ornithorhynchus anatinus (platypus, 9258). The common names and taxonomy IDs are inside the parentheses. Their untranslated regions (UTRs) were downloaded from UTRdb (32) on Nov 2024. A custom program was developed to select the longest 3’UTR isoform for each gene, resulting in 97455 (human), 57187 (mouse), 28889 (rat), 16636 (cow), and 12503 (platypus) 3’UTR sequences.

For the sequence complexity analysis (described below), human coding sequences and gene information were downloaded from the UCSC Genome Browser (33) on Nov 2024. A customized program was used to tag each 3’UTR sequence with the official gene symbol.

### Homology Information

Gene homology information was obtained from the NCBI Gene ftp site: https://ftp.ncbi.nlm.nih.gov/gene/DATA/. Two files, gene_info.gz and gene_orthologs.gz, were downloaded on May 2025. Human genes were used as the lead to identify homologous genes in the other four targeted species. A custom program was developed to select, merge, and organize gene homology information, producing 15460 homologous gene clusters in which 2902, 7349, and 5209 clusters consisted of three, four, and five species, respectively.

### 3’ UTR Alignment and Conserved Fragments Identification

MAFFT v7.525 (2024/Mar/13) (34) was used to align the 3’ UTRs in each homologous gene cluster with at least three sequences (species) using default parameters, except that the maximum iteration was set to 1000 (“--maxiterate 1000”).

A window-based approach was used to detect conserved fragments from the alignments by examining the proportion of perfect matches within a 20-bp window. In the first step, it marked all positions with perfect matches, i.e., positions where the same nucleotide occurred in all aligned sequences. In the second step, it calculated the proportion of identical matches within a 20-bp window. All positions within the window were marked as conserved if the proportion was ≥0.9. This step was repeated by moving the window one base at a time from 5’ to 3’ until the end of the alignment. After that, overlapping, marked windows were merged into longer fragments, namely conserved fragments (CFs). Using this method, human CF 50 bp or longer were detected in 2, 905 gene clusters, constituting the CF50 group. The longest CF was 624 bp. We also used two other cutoff lengths: 100 and 200 bp, to create the CF100 and CF200 groups. The CF100 group contains 760 CFs from 514 genes, whereas the CF200 group contains 123 CFs from 105 genes. See Supplementary Table S1 for the list of genes harboring CFs.

Simulated 3’UTRs were created by randomly shuffling the actual 3’UTRs within each homologous gene cluster. The simulated sequences were grouped into 2902, 7349, and 5209 clusters, consisting of three, four, and five species, respectively, the same numbers as the real gene clusters. The exact procedure used to identify CFs for real 3’UTRs was applied to simulated 3’UTRs.

## BLASTN

To expand the search for CFs beyond the five targeted mammals and non-human primates, we used human CFs with a minimum length of 100 bp (n=760) to query the NCBI RefSeq database (35) using BLASTN 2.17.0 (36). These are the non-default parameters used in the search: database: RefSeq RNA; excluded all five targeted mammals above and primates; excluded predicted sequences (XP/XM); excluded environmental samples; optimized for highly similar sequences; and turned off low-complexity. BLAST hits with a minimum coverage of 90% were retained for analysis. In total, 165 CFs were found beyond the five targeted mammals. Sus scrofa (60), Equus caballus (37), and Canis lupus familiaris (28) showed up the most. See Supplementary Table S2.

### Positional Distribution of CFs in 3’UTRs

Only human and mouse CFs were explored. Their genomic locations of CFs from the CF50 group and 3’UTRs were mapped to the human genome (hg38) and the mouse genome (mm39) using the standalone BLAT v. 39x1 (37). The outputs were kept in BED format. As the lengths of 3’UTRs vary, all 3’UTRs were standardized to a length of 100. As such, the locations of CFs were rescaled between 1 and 100, 1 being the rightmost position right after the stop codon, and 100 being the leftmost position, i.e., the 3’-end of a 3’UTR.

### Dinucleotide Composition Analysis

To reveal CF’s sequence properties, we compared CF’s dinucleotide composition with human 5’UTRs, CDS, and 3’UTRs. Human coding sequences (CDS) were downloaded from UCSC Genome Browser on Nov 2024. In total, 75, 671 CDSs were used in this study. A customized Python program was developed to break down sequences into dinucleotides.

### t-SNE Plot

t-SNE plot was produced using the R package Rtsne (38). A sequence was transformed into a vector of k-mer proportions. For instance, for k=2, a sequence is represented as a 16-dimensional vector. In this study, we created a t-SNE plot for four types of human sequences: CF50 group, CDS, 5’UTR, and 3’UTR. The number of CFs included was 893. Because the number of other sequence types was far greater than the number of CFs, 1000 CDS, 5’ UTR, and 3’ UTR sequences were randomly selected for analysis.

The maximum number of iterations was set to 1, 500 such that the error converged to a stable value. We tried different values for perplexity (5 to 50 in steps of 5) and settled on 20 for the present t-SNE plot.

### Benchmark Sequences

Additionally, we synthesized low-complexity sequences consisting of 1 (SIM1NT), 2 (SIM2NT), 3 (SIM3NT), and 4 (SIM4NT) nucleotides. In each case, nucleotides in each position are independent of their neighboring nucleotides, and the sequence length was 1000 bp.

Simulated 3’ UTR sequences were generated according to the observed nucleotide composition of the human 3’ UTRs, where A, C, G, and T occur 27.5%, 22.5%, 23.5%, and 26.5%, respectively. Each simulated 3’ UTR was 1000 bp long. They were grouped in a manner that mimicked the real gene clusters, producing 5, 363, 7, 354, and 2, 808 clusters that consisted of five, four, and three species, respectively. In total, 15, 525 fictitious clusters were created, the same number as the real gene clusters.

### Complexity of DNA Sequences

Byte Pair Encoding (BPE) was used to assess the complexity of human CDSs, 5’ UTRs, 3’ UTRs, and simulated sequences. Briefly, the input sequences contain only four letters: ‘A’, ‘C’, ‘G’, and ‘T’. BPE starts by substituting the most frequent nucleotide pair (dinucleotide) in the input sequences with an unused letter. For example, if ‘TT’ is found to occur the most, then all ‘TT’s are substituted with an unused letter, say ‘B’. Such substitution is recorded and reported when the process that necessitates decompression finishes. The substitution step will repeat until either the unused letters are exhausted or a preset number of rounds of substitution has been reached (the option we chose). The compression ratio is defined as: 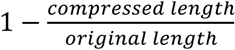 BPE was applied to SIM1NT, SIM2NT, SIM3NT, SIM4NT, CF50, CF100, CF200, CDSs (n=19439), 5’ UTRs (n=15187), and 3’ UTRs (n=18949). The minimum length of sequences was 100 bp. Note that the start and stop codons of CDSs were removed before BPE processing.

### RepeatMasker

We used RepeatMasker (Galaxy Version 4.1.5+galaxy0) (39) on the Galaxy platform (40) to examine low-complexity regions and repeats using the DFam database (41) for the human species. 2905 human CFs from the CF50 group (259504 bp) were scanned. For comparison purposes, 2000 random samples of CDSs (3579255 bp) and 2000 random samples of 3’UTRs (10463 bp). See Supplementary Table S4.

## GROVER

The GROVE pre-trained model was installed from here: https://huggingface.co/PoetschLab/GROVER. Our model was developed on a GPU server with GPU Tesla T4, and CUDA version 12.7. These additional Python libraries were installed:

captum==0.8.0

peft==0.17.1

scikit-learn==1.7.2

scipy==1.15.3

torch==2.6.0+cu124

torchaudio==2.6.0+cu124

torchmetrics==1.8.2

torchvision==0.21.0+cu124

transformers==4.57.1

Positive samples were taken from the CF50 group, which contains 2, 989 human CFs. For the negative samples, 3’UTRs confirmed with no CFs were randomly selected, resulting in 5, 203 sequences. Importantly, the lengths of the negative sequences were maintained the same as those of the positive samples. For example, if the length of a positive sequence is 213 bp, we splice a 213-bp 3’UTR segment from a random 3’UTR, starting at a random position. As recommended by GROVER, 100 bp were padded at the beginning and end of both positive and negative sequences. It was done by mapping the genomic coordinates of CFs to hg38 using standalone BLAT v. 39x1, and the output .BED file was uploaded to the UCSC Genome Browser [cite] as a custom track. Then, used the menu options Tools/Table Browser/Choose the newly created custom track/Download sequence to add 100 bp before and after the sequence.

During training, the positive and negative datasets were balanced by randomly selecting 2, 989 sequences from the negative training set. A 5-fold cross-validation was used to construct the model. Below are the hyperparameters used:

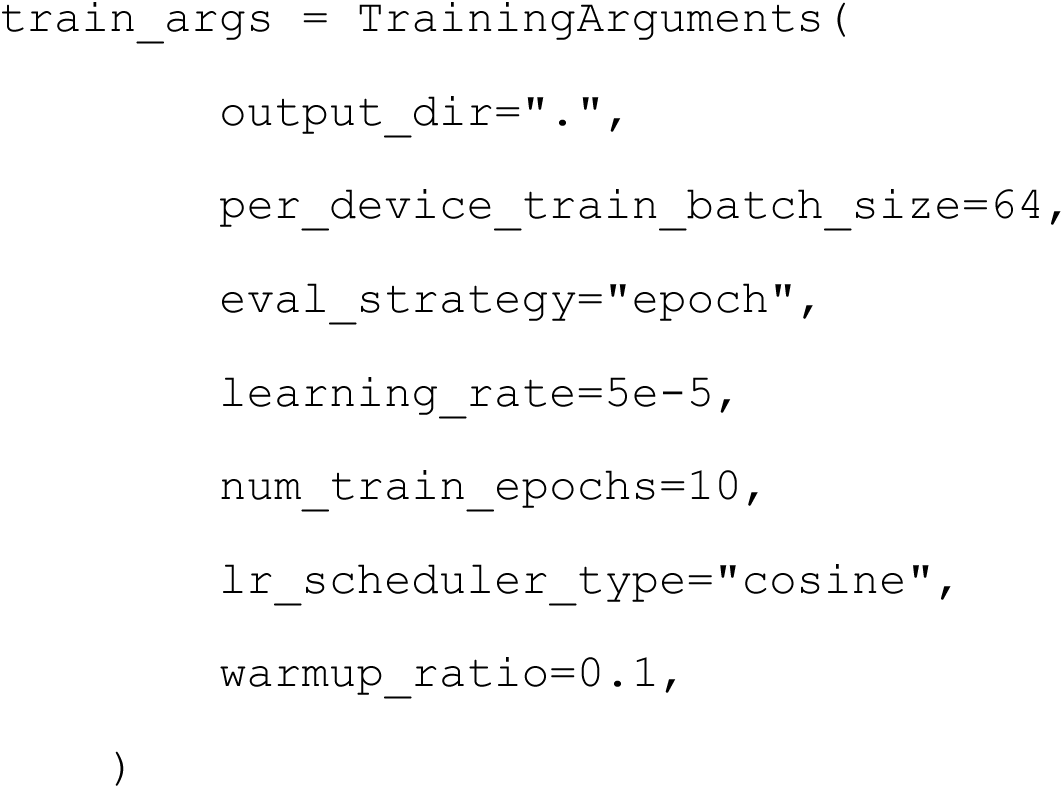

LoRA was used for fine-tuning the model (42). Below are the parameters:

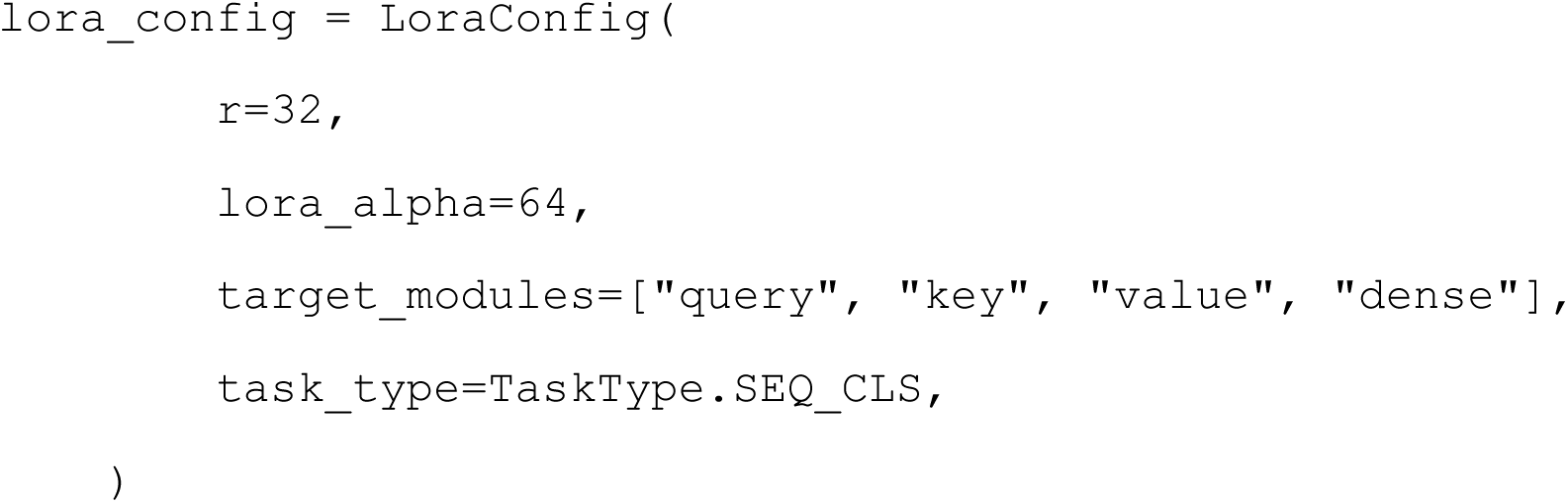

The model’s accuracy was evaluated using F1, Matthews Correlation Coefficient (MCC), Precision, and Recall. They are defined below:

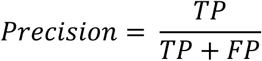

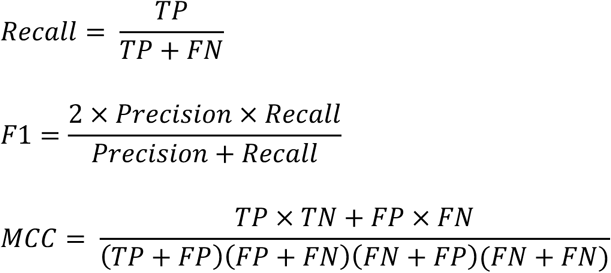

Where TP is true positive, FP is false positive, TN is true negative, and FN is false negative. The metrics’ statistics were collected by repeating the training step 10 times.

### Integrated Gradients

Explainability analysis was performed using Integrated Gradients (43). Since we trained 10 models above, the average attribution for each token (n=601) was calculated for each model. We then selected the top 10 tokens with the highest median attribution across the 10 models. See Supplementary Table S5.

### Identification of Top Tokens’ Neighbors

A custom Python program was developed for this task. First, it loaded a trained/fine-tuned CF classification model and the tokenizer. Next, it retrieved the BERT embedding via the model. And then, it processed the top 15 tokens identified by Integrated Gradients mentioned above one by one. For each token, it used the torch.topk() function to determine the k closest neighbors, where k=5. As such, 64 tokens were identified in total: 15 are the top tokens, and 49 are their neighbors. Note that some top tokens are also neighbors of other top tokens. See Supplementary Table S6.

### Finding Tokens in Training Samples

Human CF50 sequences (n=2989) and segments of human 3’UTRs with no CFs (n=5203) were selected if they achieved a prediction score greater than or equal to 0.8. A custom Python program was developed to tokenize these sequences using the same tokenizer associated with the trained model. The 64 tokens were tallied across the tokenized sequences. Refer to Supplementary Table S7 for the counts and proportions.

### Proportion Test

The R prop.test() function was used to test whether the proportion of the 64 tokens is the same between the CFs and non-CFs samples used to train the GROVER model. Recall that the number of CFs and non-CFs samples is 2989 and 5203, respectively. The counts, proportions, p-values, and -log10 p-values are provided in Supplementary Table S7.

### Constraint Metrics

Two distinct constraint metrics were used in this study: JARVIS (40) and UK Biobank depletion rank score (DR) (45). Both metrics were downloaded from UCSC Genome Browser for the human genome hg38 on Dec 2025. They were converted from bigWig to bedgraph using the UCSC GB tool bigWigToBedgraph. Next, they were mapped with the genomic coordinates of CFs from the CF50 group, and the median score was calculated for each CF. A custom Python program was developed to select intolerant CFs with a JARVIS score of at least 0.9998 and a DR of no more than 0.1, yielding 36 intolerant CFs across 25 genes. They are listed in Supplementary Table S8.

### Mapping RNA-binding Proteins Binding Sites

RNA-binding protein (RBP) binding results from eCLIP experiments were downloaded from ENCODE (46) in July 2025. In total, there are 250 RBP datasets, of which 145 are from K562, and 105 are from HepG2. The downloaded binding results were already mapped to the human genome hg38. To identify RBPs that bind to CFs, we mapped CFs to the human genome hg38 using standalone BLAT v. 39x1. And then, bedtools intersect (47) was used to determine genomic coordinates overlapping between RBPs and intolerant CFs. The -F option was 1.0 to ensure that the entire RBP binding site falls within the CF. See Supplementary Table S8. Table for details.

### Mapping Human siRNAs

Genome (hg19) locations of human predicted (conserved) targets of conserved miRNA families were downloaded from https://www.targetscan.org/cgi-bin/targetscan/data_download.vert80.cgi on Dec 2025. The UCSC Genome Browser Liftover tool was used to translate genomic coordinates from hg19 to hg38. Bedtools intersect was used to determine genomic coordinates overlapping between siRNA and intolerant CFs. The -F option was set to 1.0 to ensure that the entire siRNA-binding site falls within the CF. See Supplementary Table S8. Table for details.

### Functional Enrichment Analysis

R Enrichr package (48) was used for enrichment analysis. Enrichr analysis requires setting up background genes. We chose human protein-coding genes as the background genes. All human gene symbols were downloaded from here: https://ftp.ncbi.nlm.nih.gov/gene/DATA/GENE_INFO/Mammalia/Homo_sapiens.gene_info.gz on Sep 2025.

As a result, the background contained 19473 human genes. The general process involved selecting databases based on the category of interest. In our case, we focused on tissue enrichment, GO Process enrichment, and GO Molecular Function enrichment. The database choices are listed in the main text. In all categories, only terms with adjusted p-value < 0.05 were considered.

### GeneAgent

We used GeneAgent web tool (49) to elucidate the concerted function of siRNAs or RBPs that bind to intolerant CFs. GeneAgent uses a Large Language Model (LLM) for gene set analysis (GSA); distinct from statistical-based GSA tools, such as GSEA (50), DAVID (51), etc. The GeneAgent website (https://www.ncbi.nlm.nih.gov/CBBresearch/Lu/Demo/GeneAgent/geneagent.html) was assessed in January 2026. In each submission, we included the gene harboring an intolerant CF and the siRNAs or RBPs that bind to it. GeneAgent processed the information in five steps: Initial generation of gene set, self-verification of the process name, modification, self-verification of the analytical narratives, and summarization. In total, 30 sets of genes and siRNAs/RBPs were submitted.

## RESULTS

### Long Conserved Fragments in 3’UTRs are Unusual

Human 3’UTRs and homologous 3’UTRs from at least two of the targeted species, i.e., mouse, rat, cow, and platypus, were grouped into clusters per gene, resulting in 17, 628 gene clusters, where each gene cluster contains 3 to 5 species. (Figure. 1A) 3’UTRs in each gene cluster were aligned by MAFFT (34), followed by percentage of identity and length filtering. Aligned fragments that exhibit 90% or more identity and a minimum length of 50 bp, 100 bp, or 200 bp were extracted into three conserved fragment (CF) groups, namely CF50, CF100, and CF200. Note that CF50 is the biggest group as it includes CFs from CF100 and CF200.

**Figure 1.**
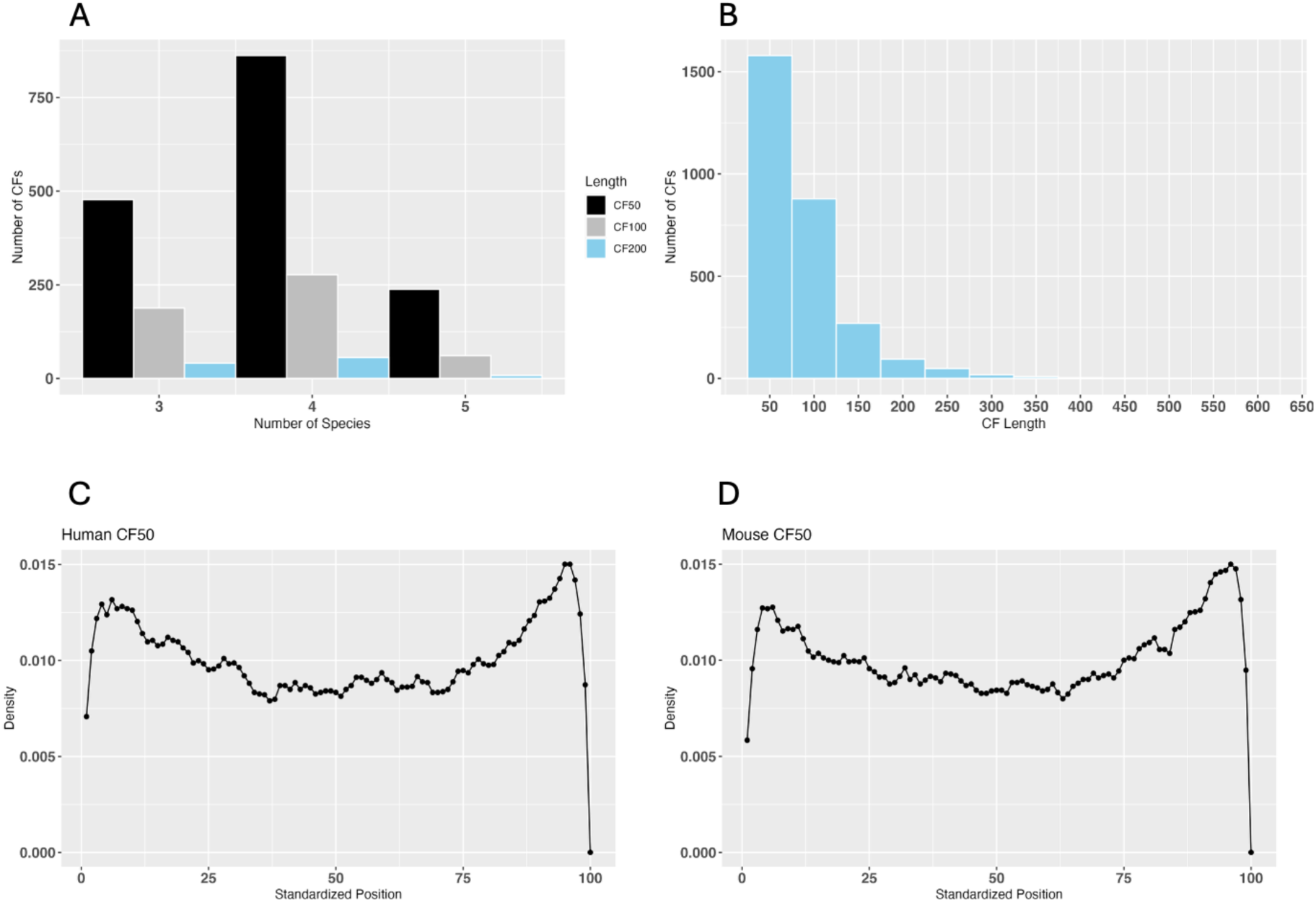
Basic statistics of conserved fragments (CFs). A. It shows the number of CFs discovered from gene clusters with three to five species. The majority of gene clusters comprises four species, i.e., human and three other targeted species. B. Distribution of CF lengths. 2905, 760, and 123 CFs are at least 50 bp, 100 bp, and 200 bp long, respectively. C. Positions where human CFs (≥50 bp) are detected with the longest 3’UTR as the reference. The length of a 3’UTR is standardized to 100, where positions 1 and 100 are the 5’-end and 3’-end of a 3’UTR, respectively. D. Same as panel C for the mouse.

Similarly, CF100 is a superset of CF200. The length distribution of the CF50 group is shown in Figure 1B.

Table 1 shows the number of genes and CFs for the three CF groups in humans. The complete list of gene symbols for the three groups is provided in Supplementary Table S1. The longest CF is 620 bp long and comes from PURA (Purine-rich element binding protein A), and shares 97% identity among human, mouse, rat, and cow. In terms of the total length of CFs per gene, RC3H2 (Ring finger and CCCH-type domains 2) has the most extended total length of 2739 bp, originating from 12 CFs found in the gene (Supplementary Table S1).

**Table 1.** Human conserved fragments statistics.

|  | CF50 | CF100 | CF200 |
| --- | --- | --- | --- |
| Number of genes | 1430 | 514 | 105 |
| Number of CFs | 2905 | 760 | 123 |
| Median length (bp) | 72 | 133 | 244 |

To eliminate the concern that the observed CFs are spurious, we applied the exact procedure for identifying CFs on simulated 3’ UTR sequences in which nucleotides of real 3’UTRs were randomly shuffled, ensuring their length and nucleotide composition were maintained. No fragments with length ≥50 bp that showed ≥90% identity were found, ascertaining that the occurrence of CFs by chance is statistically improbable.

### Prevalence of CFs Beyond the Targeted Mammals

We are wondering whether CFs are present beyond the five targeted mammals. Hence, 760 human CFs (≥100 bp) were used to query cDNAs deeper into other non-primate mammals deposited in the NCBI RefSeq RNA database (35) using BLASTN (52). Hits that covered at least 90% of the query CF were retained for analysis. In total, 165 out of 760 human CFs were identified in other mammals, where 60 CFs were found in Sus scrofa, 37 in Equus caballus, and 28 in Canis lupus familiaris. Intriguingly, the CF of two antizymes, AZIN1 (Antizyme inhibitor 1) and OAZ2 (Ornithine decarboxylase antizyme 2), were found in 47 and 36 mammals, respectively (see Supplementary Table S2). When the search was expanded to non-mammalian species, we found CFs even in chicken (Gallus gallus) and the African clawed frog (Xenopus laevis). It indicates that CFs are more widespread at the 3’ UTRs. That said, the result cannot be generalized, as sequence data from non-model organisms are limited, underscoring the crucial need for additional data.

### CFs Concentrate at 3’UTRs’ Ends

CFs are found mainly localized at the two ends of the 3’UTRs, as shown in Figure 1C-D. Such a pattern is conserved between human and mouse. A small number of them (14 pairs) are found at the extreme 3’-end of 3’UTRs, due to gene overlap. Additionally, the gene overlaps are evolutionarily conserved between human and mouse (Supplementary Table S3).

However, it is a tiny fraction of the CFs found (28 out of 2905, see Table 1 above) near the 3’-end of the 3’UTRs. Other reasons are needed to explain the large number of CFs near the 3’-end.

### CFs are AT-rich and Distinct from 5’UTR and CDS

The proportion of dinucleotides was examined across four sequence types: CDS, 5’ UTR, 3’ UTR, and CF. As CFs among the five studied mammals are highly similar by definition, only human sequences were examined here. The conclusion is expected to hold for other organisms.

It is found that AT, TA, and TT dinucleotides in CFs have medians greater than the 75% quartile of other sequence types (Figure 2A). It is worth noting that, despite AA being the highest in CFs among other sequence types, its median does not pass the 75% quartile of the other three. These observations suggest that CFs are enriched by ‘T’s, and alternating T’s and A’s, instead of A-runs. Notably, no dinucleotide over- or underrepresentation is observed in the 3’UTR sequence type (green in Figure 2A), even though they host the CFs.

**Figure 2.**
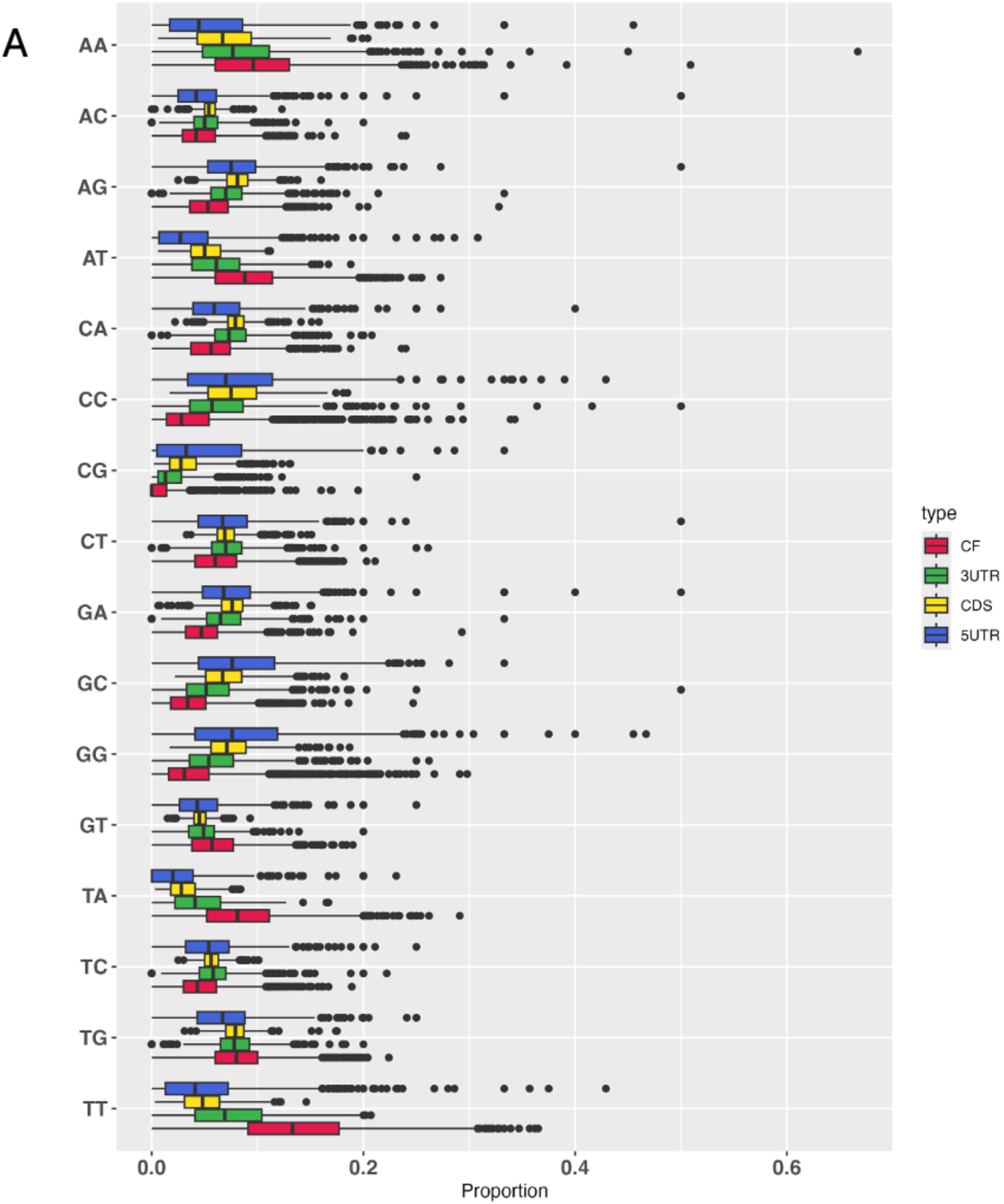
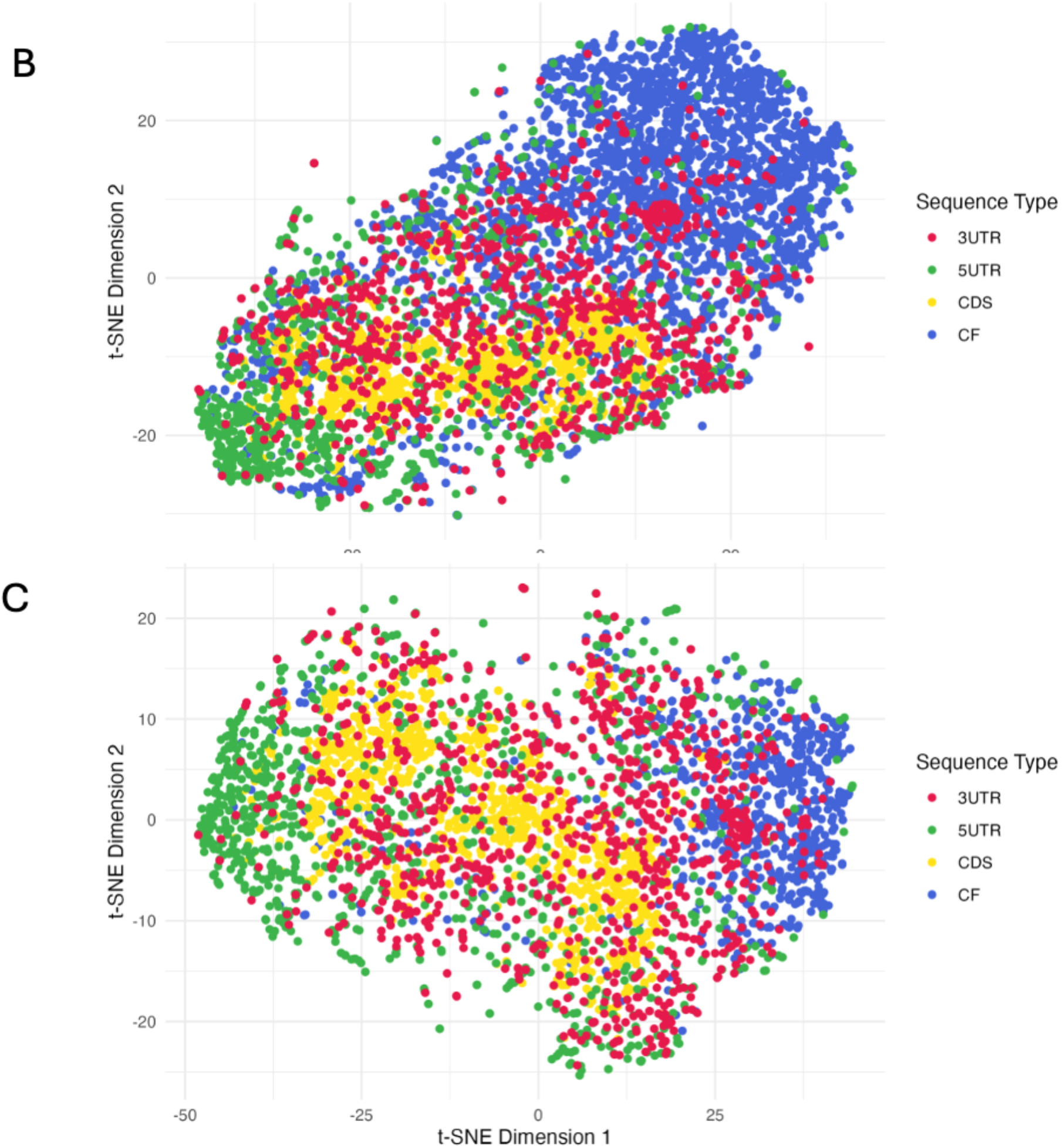
Nucleotide composition of CFs. A. Dinucleotide composition of CFs when compared with 5’UTR, CDS, and 3’UTR in humans. The median proportions of AT, TA, and TT of CFs are higher than the 75% quantile of other sequence types. B. t-SNE plot for CF50 dinucleotides. CFs are clustered and overlap little with CDSs (yellow) and 5’UTRs (green).

However, 3’UTRs (red) are scattered but distinct from 5’UTRs. C. t-SNE plot for CF100 dinucleotides. CFs are more tightly clustered than CF50, suggesting that the sequence characteristics of longer CFs are distinct from shorter CFs. The t-SNE plot for CF200 is shown in Supplementary Figure S1.

We further compared the nucleotide composition of CFs with other sequence types using the t-SNE plot. The plots for human CF50 and CF100 dinucleotides with 5’UTR, CDS, and 3’UTR are shown in Figure 2B-C. The segregation between CF50’s CFs and CDSs is noticeable with minimal overlap, even more distinct for the CF100 group. The cluster of 5’UTRs is closer to the CDS, but distant from CFs, suggesting that the nucleotide composition of CFs is distinct from 5’UTRs and CDS. However, the cluster for 3’ UTRs is closer to that of the CFs, and yet they are distinguishable. The t-SNE plot for CF200 is provided in Supplementary Figure S1.

### CFs are not Low-Complexity

Low-complexity and short repeats can lead to spurious sequence similarity. To eliminate such a possibility for CFs, we employ the idea of compressibility to assess whether CFs are composed of short repeating sequence units. Intuitively, if sequences are composed of simple repeats, they can be compressed to a greater degree than complex, non-repetitive sequences. We used the compression method Byte Pair Encoding (BPE), which is widely used in tokenizing natural language. Here, we used the compression ratio attained by BPE as a measure of the complexity (or simplicity) of CFs. We benchmarked the compression of CFs against real 5’UTR, CDS, and 3’UTR sequences. Additionally, simulated sequences that consist of one to four different nucleotides were generated, namely SIM1NT, SIM2NT, SIM3NT, and SIM4NT.

Figure 3A shows the trend in the compression ratio for various sequence types by iterations. Generally speaking, in a specific iteration, the compression ratio of a complex, non-repetitive sequence (e.g., SIM3NT and SIM4NT) is smaller than that of a sequence containing simple repeats (e.g., SIM1NT and SIM2NT). As shown, the compression ratio of the longest CF (from PURA) follows a similar trend to those of its CDS, 5’ and 3’ UTRs, mirroring that of SIM3NT. It is evident that the CF from PURA does not fill with repeats. Figure 3B compares all CFs with CDSs, 5’, 3’UTRs, and simulated sequences. The medians of CFs are slightly higher (Figure 3B, right) than those of CDSs and 5’-/3’UTRs. The median compression ratio across the CF groups remains similar. Within-group variation of CFs diminishes from CF50 to CF200, which is probably attributed to sample size and noise in shorter CFs. Nevertheless, outliers in CFs, regardless of group, are observed only at lower compression ratios, reflecting more complex sequence patterns across the three CF groups. To that end, BPE reviews that CFs are not dominated by simple repeats.

**Figure 3.**
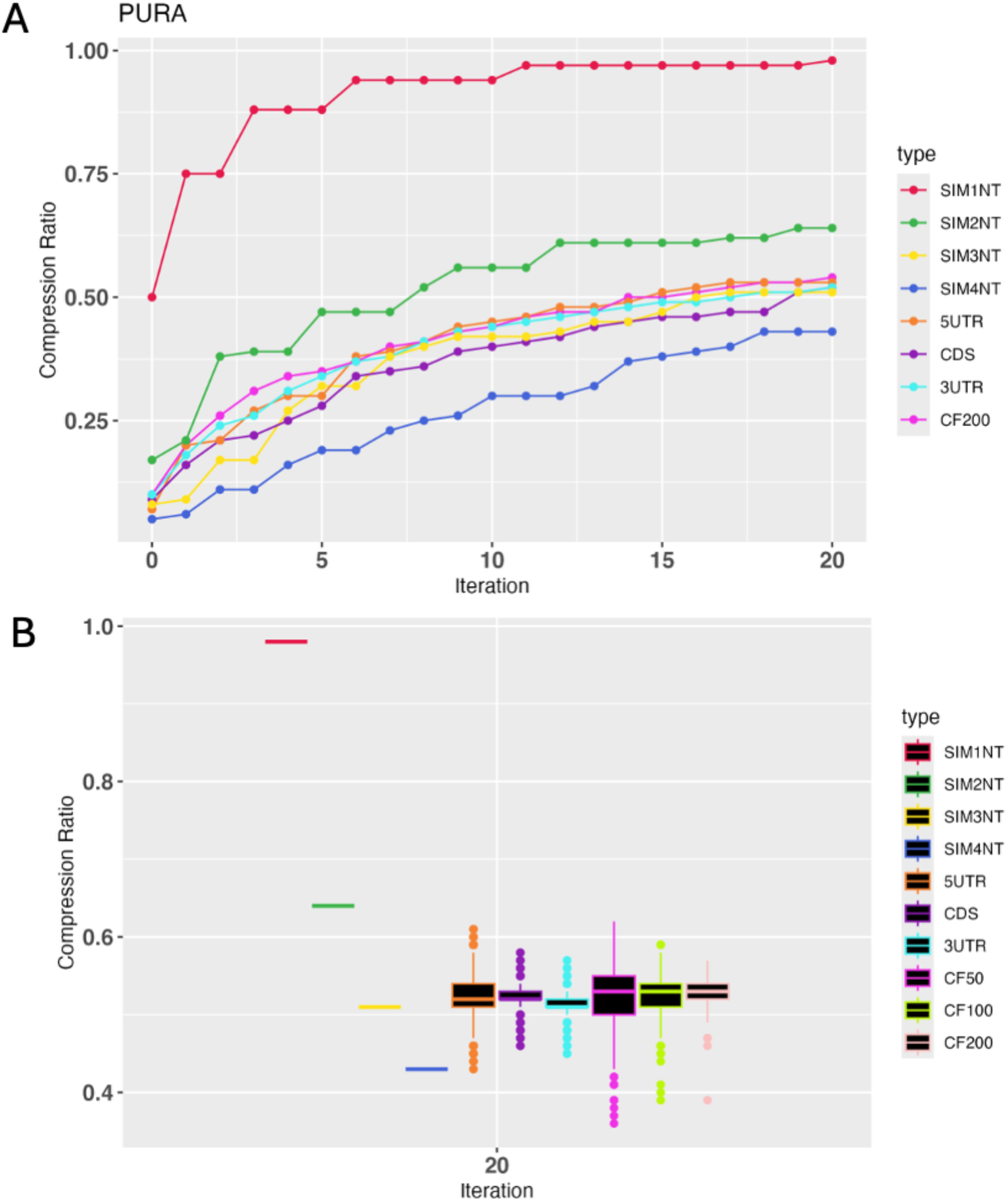
Low-complexity assessment by BPE. A. An example of a particular gene PURA. The degree of compression of the real sequences (5’UTR, CDS, 3’UTR, and CF of PURA) by iteration is similar to a simulated sequence consisting of three different nucleotides (SIM3NT). B. Robust statistics of compression ratio for all sequences. The overall degree of compression of CFs from the three groups is similar to 5’UTR, CDS, and 3’UTR. Among the CFs, within-group variation reduces from the CF50 to the CF200 group, partly due to the sample size. Importantly, outliers are located below the lower whisker, indicating complex sequence patterns.

To corroborate the compressibility analysis, we wanted to rule out the possibility that CFs are inundated with low-complexity repeats, which constitute 50% of the human genome (53).

RepeatMasker (39) was used to detect simple repeats and low-complexity sequences in CFs (based on the CF50 group), CDSs, and 3’UTRs. In all categories (SINE, LINE, LTR elements, DNA elements, unclassified, and interspersed repeats), 3’UTR contains the highest percentage compared to CDS and CF (see Supplementary Table S4). More importantly, no LTR retrotransposons were found in CFs, eliminating the viral origin factor. CFs carry no satellites and have as few as 0.05% of low-complexity repeats, smaller than CDSs.

### CFs under the Lens of Transformer

At this point, we have learned that CFs are enriched with TT, TA, and AT dinucleotides. However, they are not saturated with low-complexity elements that are ubiquitous in the human genome. We are wondering what regulatory signals are preserved in the CFs over millions of years of evolution. Because their biological functions remain unknown and their lengths are much longer than typical sequence motifs, such as transcription factor binding sites, existing motif finders are not suitable for this task. Thus, we harnessed the DNA foundation language model GROVER (31) to unearth the sequence characteristics associated with the CFs. GROVER used the BPE mentioned above to tokenize the human genome into 601 tokens. Each token is a k-mer, where k ranges from 1 to 16. The shortest tokens of size 1 are individual nucleotides, and the longest tokens are a run of 16 A’s and Ts.

By using transfer learning, we built a binary classifier using the pre-trained GROVER model to distinguish CFs (n=2989) from 3’UTRs that possess no CFs (n=5203). The use of 3’UTRs as the negative samples is justified, as CFs originate from 3’UTRs. Factors that specifically target CFs must distinguish between the two for specific interactions. To avoid issues with imbalanced data, the number of negative samples was kept equal to that of the positive samples via random sampling. Five-fold cross-validation was used during training. Evaluation statistics were obtained by training the model repeatedly 10 times. The model’s accuracy is assessed by F1, Matthews Correlation Coefficient (MCC), Precision, and Recall. The median F1, MCC, Precision, and Recall are 0.8329, 0.6734, 0.8396, and 0.8341, respectively. (Figure 4)

**Figure 4.**
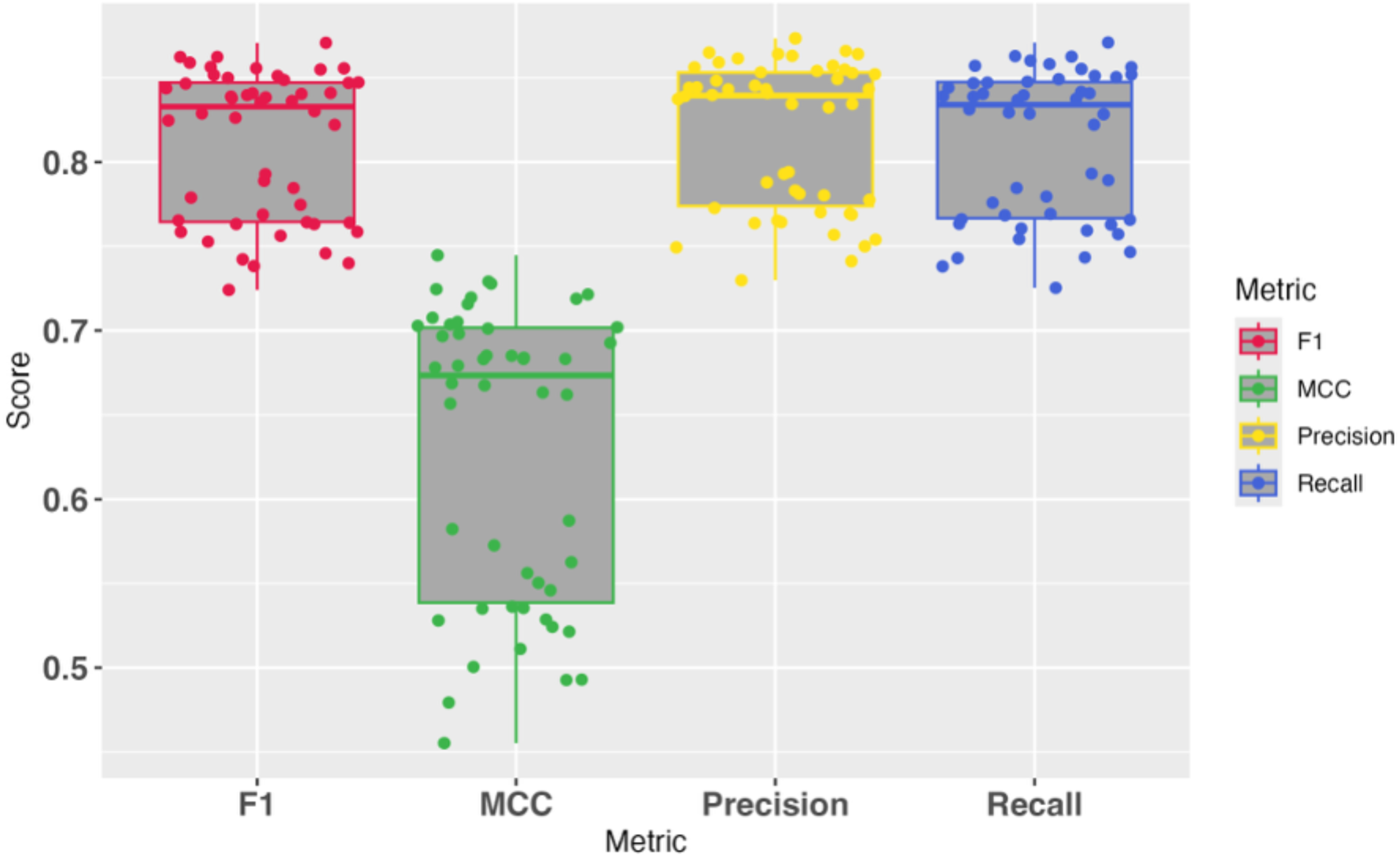
The accuracy of the GROVER-based CF model. The model was trained using 5-fold cross-validation repeated 10 times. The median F1, MCC, Precision, and Recall are 0.8329, 0.6734, 0.8396, and 0.8341, respectively. Each dot represents a metric value from a test run. The ten repeats of 5-fold cross-validation The model has successfully distinguished the two sequence types and achieved F1, precision, and recall greater than 0.8. However, a considerable variation is observed in the MCC. The gap between F1 and MCC is attributed to two factors. First, F1 primarily focuses on predicting positives (i.e., CFs), while MCC integrates both positive and negative predictions (i.e., 3’UTRs with no CFs). It is related to the second factor, in which 3’UTRs are diverse. It is known that 3’UTRs are subjected to a weaker selective pressure than CDSs, for instance. Their diversity is also evident in the t-SNE plots in Figure 2B and 2C. The heterogeneous nature of the negative samples, which are randomly sampled from a much larger 3’UTR space. That poses a challenge for the model in confidently predicting negative samples. Despite this, our objective is to identify signatory sequences embedded in CFs, as F1, precision, and recall are all above 0.8, suggesting that the model has captured sequence characteristics that adhere to CFs. Lastly, the correlation between CF length and prediction scores was examined. It found no correlation (r=0.02).

### Characteristics of CFs

Next, we set out to identify distinct sequence patterns captured by the model that differentiate CFs from non-CFs. The explainability method Integrated Gradients (43) was used to tap into the model parameters and score the 601 tokens of GROVER that contribute to its predictions, namely attribution. The average and median attributions of each token across the 10 repeated training runs were calculated and ranked (see Supplementary Table S5).

The 15 tokens with the highest median attribution are colored in blue dots in Figure 5A. To understand sequence variability, we also incorporated the five closest neighboring tokens of these 15 top tokens in GROVER’s embedding space, based on cosine similarity, and colored them in grey dots (Figure 5A). In total, 64 tokens are considered. The thickness of edges represents cosine similarity (see Supplementary Table S6).

**Figure 5.**
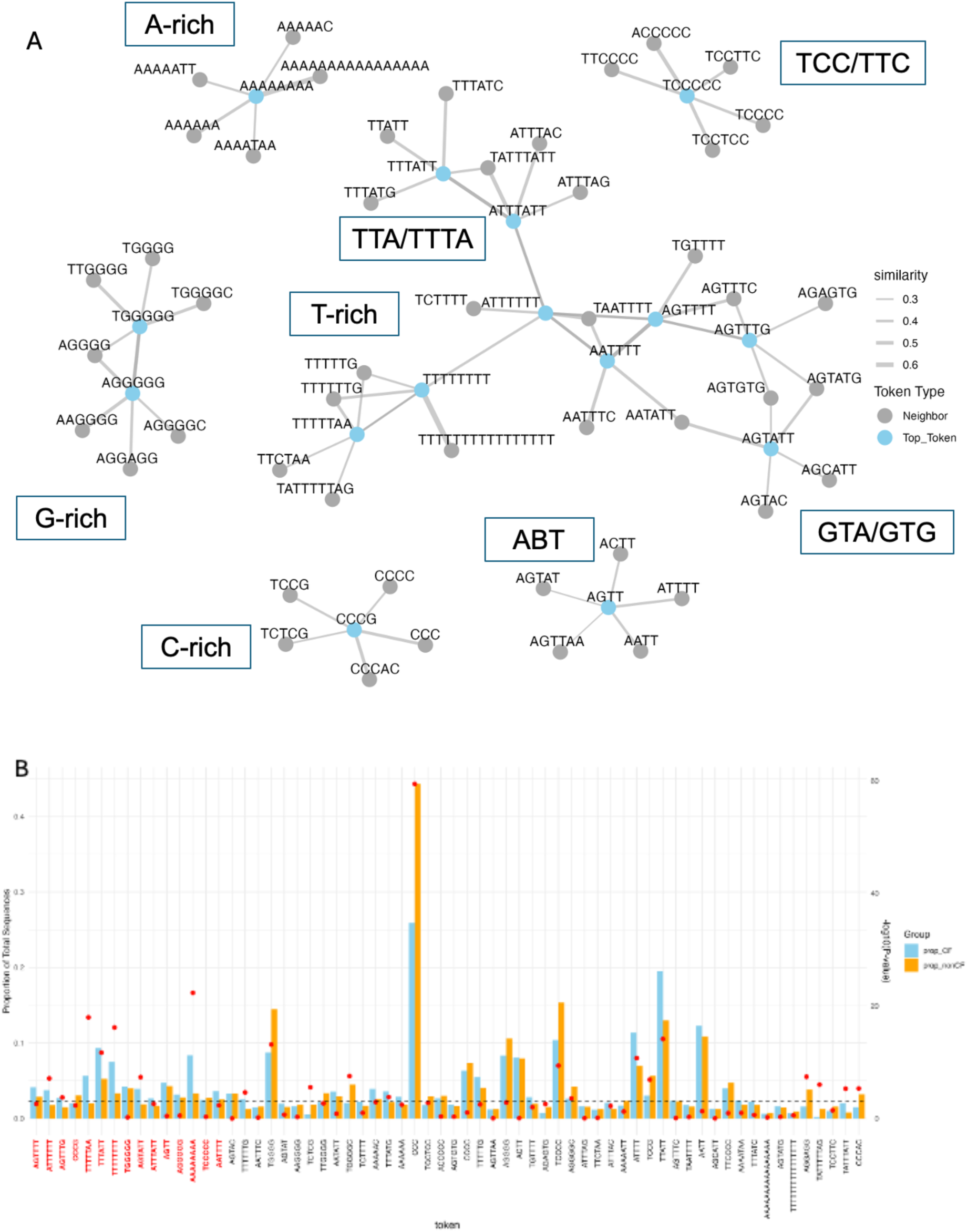
Explainability analysis. A. A network of top 15 tokens (blue) identified by Integrated Gradients. The top tokens are connected to their five nearest neighbors (grey) based on GROVER’s embedding. The thickness of the edges represents cosine similarity. There are 64 tokens in total. Note that some top tokens are also neighbors of other top tokens, e.g., ATTTTTT and ATTTATT are top tokens and neighbors. B. The proportion of CFs contains a token. The total number of CFs and non-CFs is 2989 and 5203, respectively. Top Tokens are highlighted in red on the left. The other tokens are neighbors (black). The red dots represent the –log10 of p-values from proportion tests. The horizontal dashed line represents the adjusted p-value after Bonferroni’s adjustment, i.e., -log10(0.05/64), which is equivalent to 3.1.

These tokens form into six clusters, indicating six distinct patterns are found in CFs. The biggest one is being T-rich, in which T is the dominant nucleotide across all tokens, ranging from 1/3 to 100%. Ts are either punctuated or continuous. There is also a small cluster labelled ABT, where B means not A (i.e., G, C, or T) according to the IUPAC nomenclature. All tokens are shorter (4-5 bp) than other clusters, begin with A, and almost all contain two Ts, except for ATTTT. The A-rich and G-rich clusters constitute clearly homopolymeric A and G. Lastly, the C-rich and TCC/TTC clusters are enriched with a run of 2 to 4 Cs. Overall, these clusters provide critical information to understand the characteristics of CFs. Importantly, these 64 tokens, out of 601, provide specific patterns that can guide downstream analyses.

Moreover, we identified the proportions of positive and negative training sequences (including 100 bp padding) that contain the 64 tokens (Figure 5B). Only sequences that are reliably predicted (prediction score ≥0.8) were selected and then tokenized. The top 15 tokens identified by Integrated Gradients above are highlighted in red on the left; the rest are their neighbors. A proportion test was used to test the null hypothesis that tokens occur at the same level in CFs and non-CFs. The -log10 p-values are displayed for each token (red dots). Bonferroni’s adjustment was used to correct for multiple testing. The cutoff adjusted p-value for statistical significance is 0.05/64 or 3.1 after taking -log10 (horizontal dashed line in Figure 5B). Tokens with p-value above the cutoff among CFs and non-CFs are tabulated in Table 3.

**Table 3.** Statistically Significant Tokens.

| CF |  | NonCF |  |
| --- | --- | --- | --- |
| Token | p-value | Token | p-value |
| ATTTTTT | 7.852089e-08 | TGGGG | 7.614483e-14 |
| AGTTTG | 1.907182e-04 | TCTCG | 2.896057e-06 |
| TTTTTAA | 1.126965e-18 | TGGGGC | 3.078462e-08 |
| TTTATT | 2.088561e-12 | CCC | 3.793659e-60 |
| TTTTTTTT | 7.045940e-17 | TCCCC | 4.174983e-10 |
| AGTATT | 4.548119e-08 | AGGGGC | 2.826885e-04 |
| AAAAAAAA | 4.731336e-23 | TCCG | 1.310195e-07 |
| TTTTTTG | 2.413110e-05 | AGGAGG | 3.934964e-08 |
| TTTATG | 1.606906e-04 | TATTTTTAG | 9.887097e-07 |
| ATTTT | 1.908592e-11 | CCCAC | 4.846523e-06 |
| TTATT | 7.763828e-11 |  |  |
| TATTTATT | 5.093050e-06 |  |  |

In consistent with the analysis above, T-rich, A-rich, and AT-rich tokens dominate CFs. On the other hand, tokens from the non-CFs are C-rich and G-rich.

### CFs Intolerant of Variations

Although the evolutionary approach we used, based on multiple sequence alignment, supports significant conservation, it remains unclear whether CFs are intolerant of genetic variation in humans. To address this question, we employ two independent metrics to assess the extent of variation constraint or depletion in human CFs. The two metrics are JARVIS (44) and depletion rank score (DR) (45). Both are based on a large number of whole-genome sequences from healthy cohorts (62784 for JARVIS, 150119 for DR). The JARVIS score ranges from 0 to 1. 0 is perfectly tolerant of variation, and 1 is totally intolerant of variation. To identify intolerant regions, the recommended JARVIS threshold is 0.9998, corresponding to the 99^th^ percentile. DR is also in the range between 0 and 1. However, 0 indicates the greatest depletion of variation, and 1 indicates the complete depletion of variation. The recommended threshold for detecting intolerant regions is ≤ 0.1.

We applied these two independently established metrics to screen for intolerant CFs from the CF50 groups (n=2989). JARVIS and DR scores were downloaded from UCSC GB for human hg38. The median score is determined for each CF, and their distribution is as shown in Figure 6.

**Figure 6.**
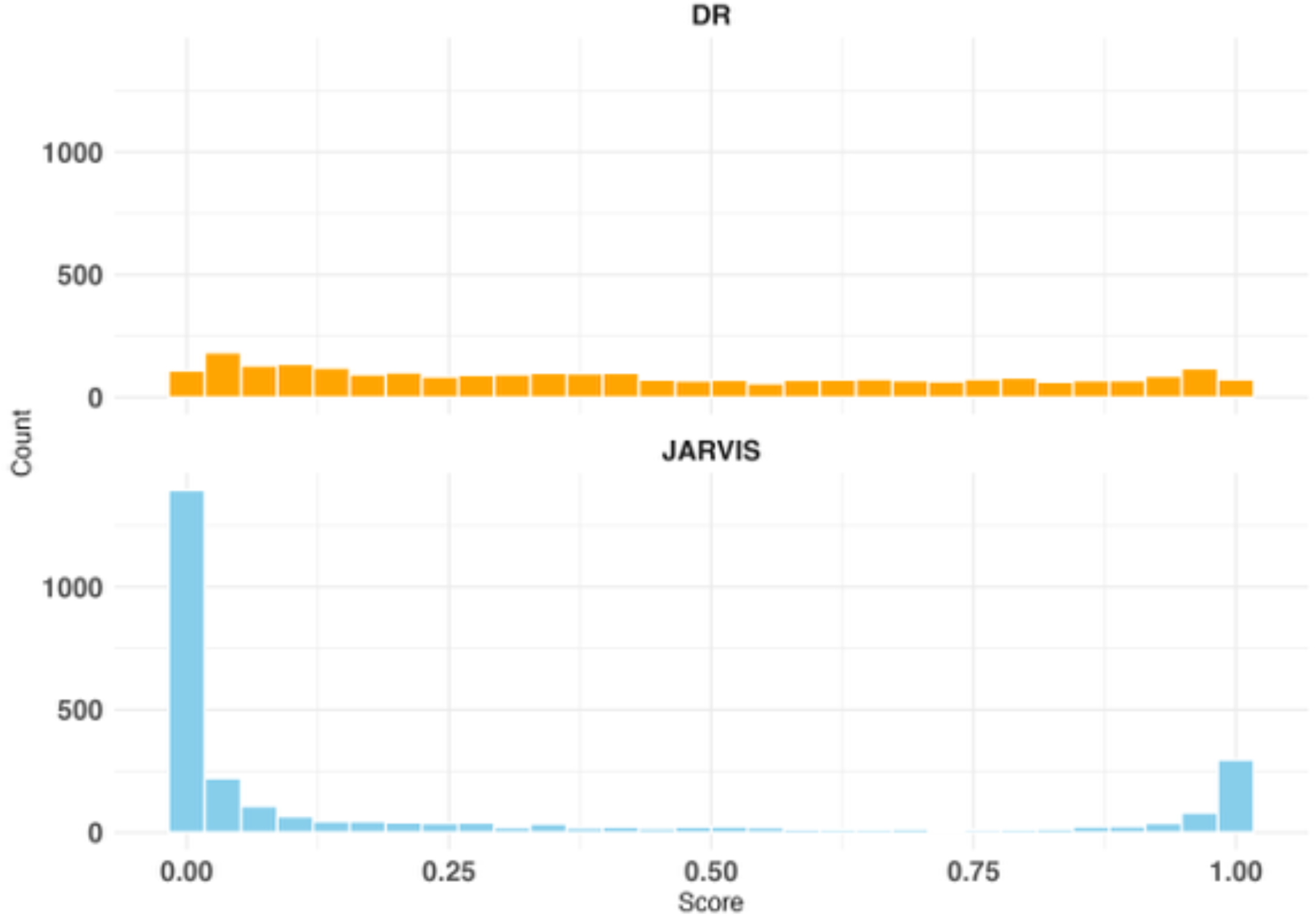
The median UK Biobank depletion rank scores (DR) and JARVIS scores of CFs from the CF50 group (n=2989). For the DR metric, scores ≤0.1 indicate depleted variation, whereas JARVIS scores ≥ 0.9998 indicate intolerant variation.

Next, we identify intolerant CFs with a JARVIS score of at least 0.9998 and a DR of no more than 0.1. As a result, 36 CFs from 25 genes were identified (Supplementary Table S8). We will name these intolerant CFs and intolerant genes.

### Functional Analysis of Intolerant Genes

We explored the shared tissue-specific expression, biological processes, and molecular functions of genes in which intolerant CFs are found. The R package Enrichr was used for the analysis (48). We scanned all library names in Enrichr with the keyword ‘tissue’, resulting in 10. But only four produced positive enrichment results: ARCHS4_Tissues, Jensen_TISSUES, Descartes_Cell_Types_and_Tissue_2021, Tissue_Protein_Expression_from_Human_Proteome_Map, and GTEx_Tissue_Expression_Up. The proteome map is based on mass spectrometry, and the rest is from RNA-Seq data. They are high-quality, comprehensive databases designed for tissue-specific analysis. Importantly, they are maintained by independent research groups, and as such, aggregated results can be used to validate significant findings. Enriched tissues with the adjusted p-value ≤0.05 are listed in Table 6. Supplementary Table S9 provides the complete list of enriched tissues with more details, including tissue databases, p-values, adjusted p-values, and intolerant CF genes that contributed to the enrichment.

**Table 6.**
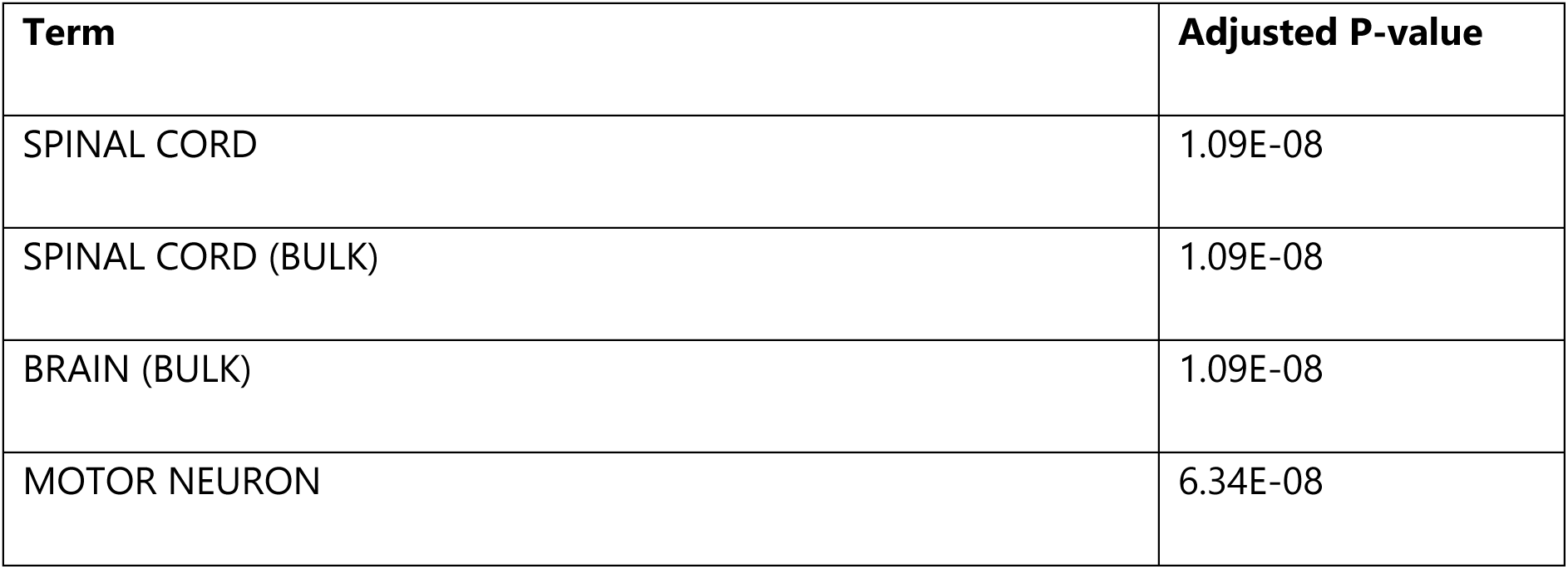
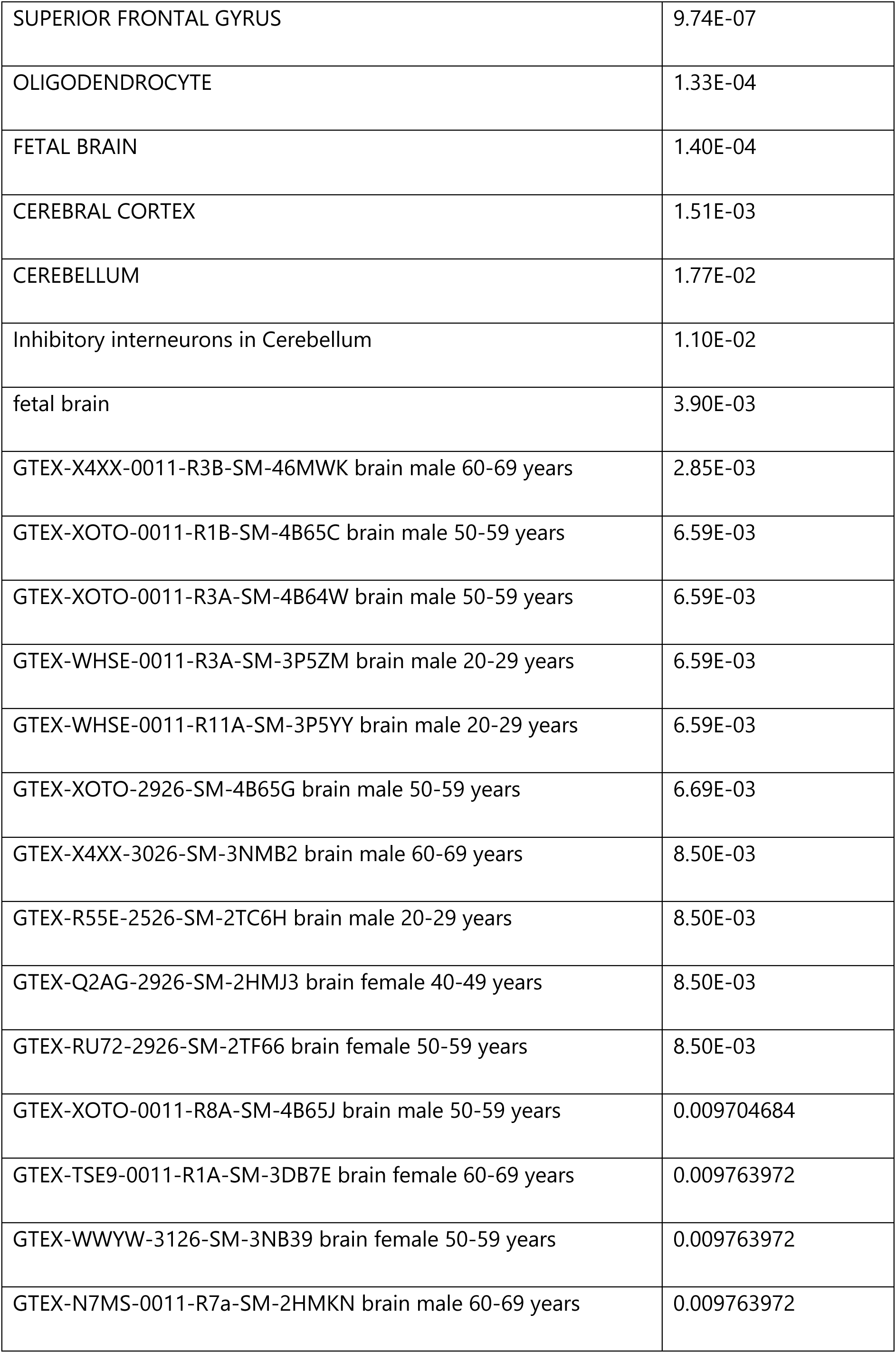
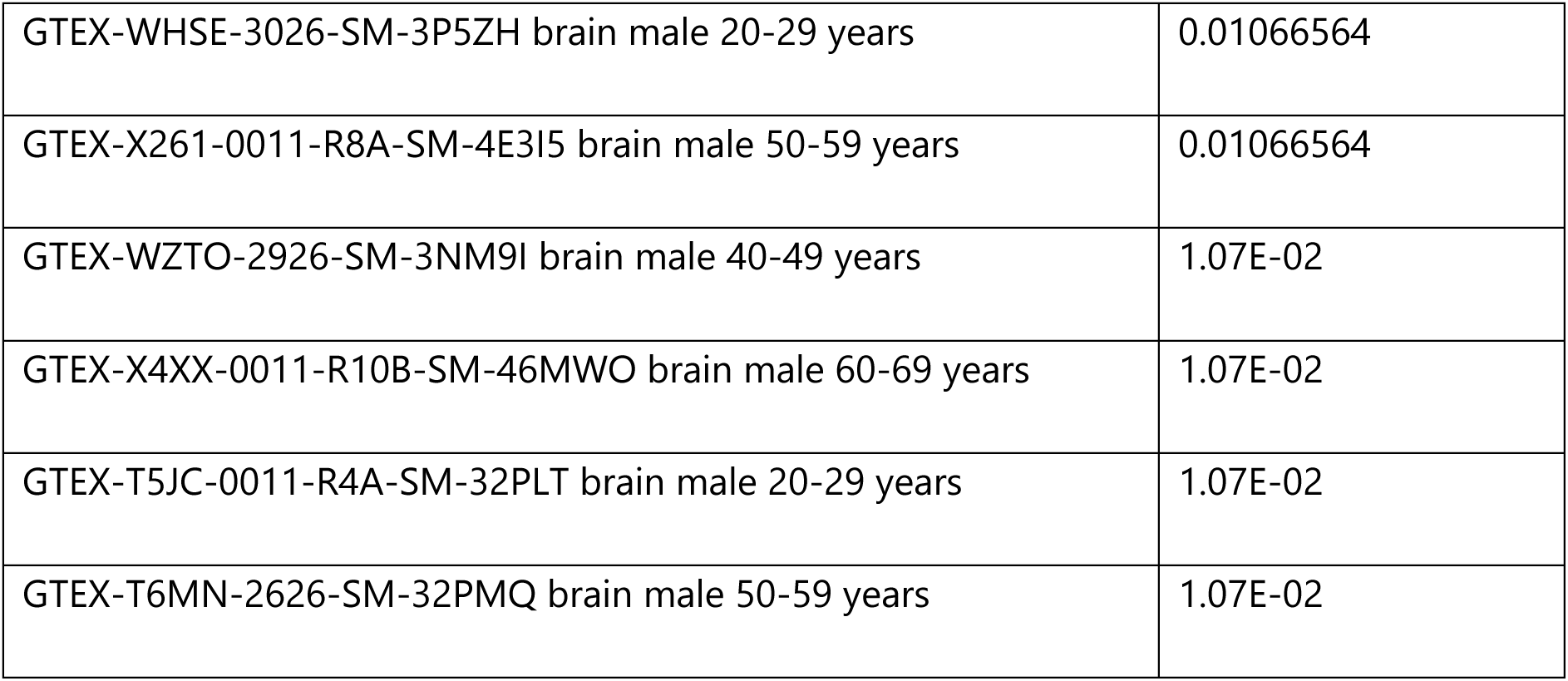
Tissue enrichment of intolerant genes sorted by adjusted p-value.

Intriguingly, all 31 tissues or samples are related to the central nervous system, suggesting probable roles of the intolerant CFs in neurons.

Next, we performed a GO process enrichment analysis using the GO_Biological_Process_2025 database. Similarly, GO processes with an adjusted p-value ≤0.05 are selected for analysis (Table 7). The results are generally split into four categories: CNS-related processes, transcriptional regulation, developmental process (GO:0048754 and GO:0048863), and kinase activity (Table 7). In particular, the adjusted p-values for the top two processes (brain development and brain morphogenesis) are the smallest, consistent with the tissue enrichment outcomes above. Moreover, kinase activities are vital regulatory processes for cell growth, metabolism, signaling, and division, and they are coupled to neuronal development.

**Table 7.** GO Process Enrichment

| **Term** | **Adjusted P-value** |
| --- | --- |
| Brain Development (GO:0007420) | 0.000590876 |
| Brain Morphogenesis (GO:0048854) | 0.007591029 |
| Camera-Type Eye Development (GO:0043010) | 0.045064685 |
| Regulation of DNA-templated Transcription (GO:0006355) | 0.045064685 |
| Branching Morphogenesis of an Epithelial Tube (GO:0048754) | 0.045064685 |
| Regulation of Transcription by RNA Polymerase II (GO:0006357) | 0.045064685 |
| Neuron Differentiation (GO:0030182) | 0.045064685 |
| Regulation of Cyclin-Dependent Protein Serine/Threonine Kinase Activity (GO:0000079) | 0.045064685 |
| Stem Cell Differentiation (GO:0048863) | 0.045064685 |
| Positive Regulation of Kinase Activity (GO:0033674) | 0.045064685 |
| Generation of Neurons (GO:0048699) | 0.045064685 |

Lastly, we investigated molecular function enrichment using the GO_Molecular_Function_2025 database. Table 8 below shows enriched molecular functions grouped into two categories: double-stranded DNA binding and kinase binding. These results align with the GO process enrichment.

**Table 8.** GO Molecular Function Enrichment

| **Term** | **Adjusted P-value** |
| --- | --- |
| Double-Stranded DNA Binding (GO:0003690) | 0.029901731 |
| Sequence-Specific Double-Stranded DNA Binding (GO:1990837) | 0.029901731 |
| Sequence-Specific DNA Binding (GO:0043565) | 0.029901731 |
| Kinase Binding (GO:0019900) | 0.033299551 |
| Protein Kinase Binding (GO:0019901) | 0.039600239 |

In summary, the enrichment analysis provides compelling evidence for a connection between genes harboring intolerant CFs and molecular processes involving the central nervous system. Potential processes encompass transcription regulation and kinase activity. Next, we will investigate factors that may deliver functions upon binding with the intolerant CFs.

### RNA-binding Proteins and siRNAs Bind to Intolerant CFs

We are curious whether intolerant CFs are targets of RNA-binding proteins (RBPs) and siRNAs, as they are known to target 3’UTRs. We incorporated RNA-binding results from eCLIP experiments downloaded from the ENCODE Project (46), which included 250 RBPs tested on the K562 (n=145) and HepG2 (n=105) cell lines. Custom Python programs and bedtools (47) were used to map RBP binding sites and the 36 intolerant CFs to the human genome (hg38). In addition to RBPs, miRNAs are key regulators of gene expression by destabilizing mRNAs, and repressing protein translation. These functions are exerted through the binding to 3’UTRs. Hence, human predicted targets of conserved miRNA families from TargetScan (54) were mapped to the human hg38 genome. As a result, 25 out of 36 intolerant CFs are found to bind to siRNAs or RBPs or both (Table 9). Some genes appear more than once in the table because they harbor multiple intolerant CFs. More details, including CF’s absence of RBPs and siRNAs bindings, can be found in Supplementary Table S9. To determine the biological function of these interactions, we used a Large Language Model (LLM) GeneAgent to elucidate the interactions between the gene harboring the intolerant CF and siRNAs or RBPs. The primary reason we used GeneAgent rather than conventional gene set analysis tools, such as Enrichr, is that they lack the novel relationship between intolerant CFs, siRNAs, and RBPs. Results are summarized in Table 9.

**Table 9.**
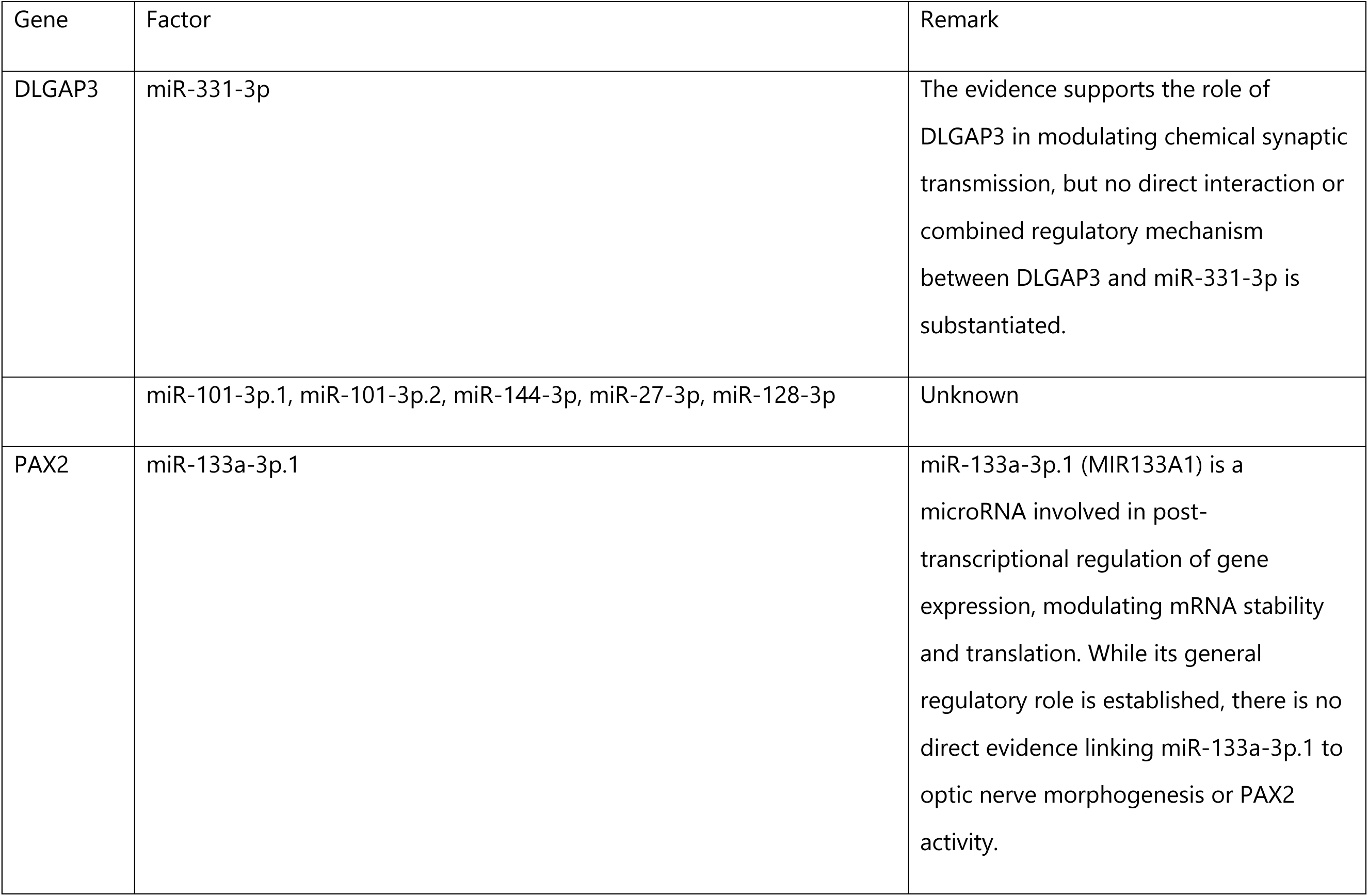
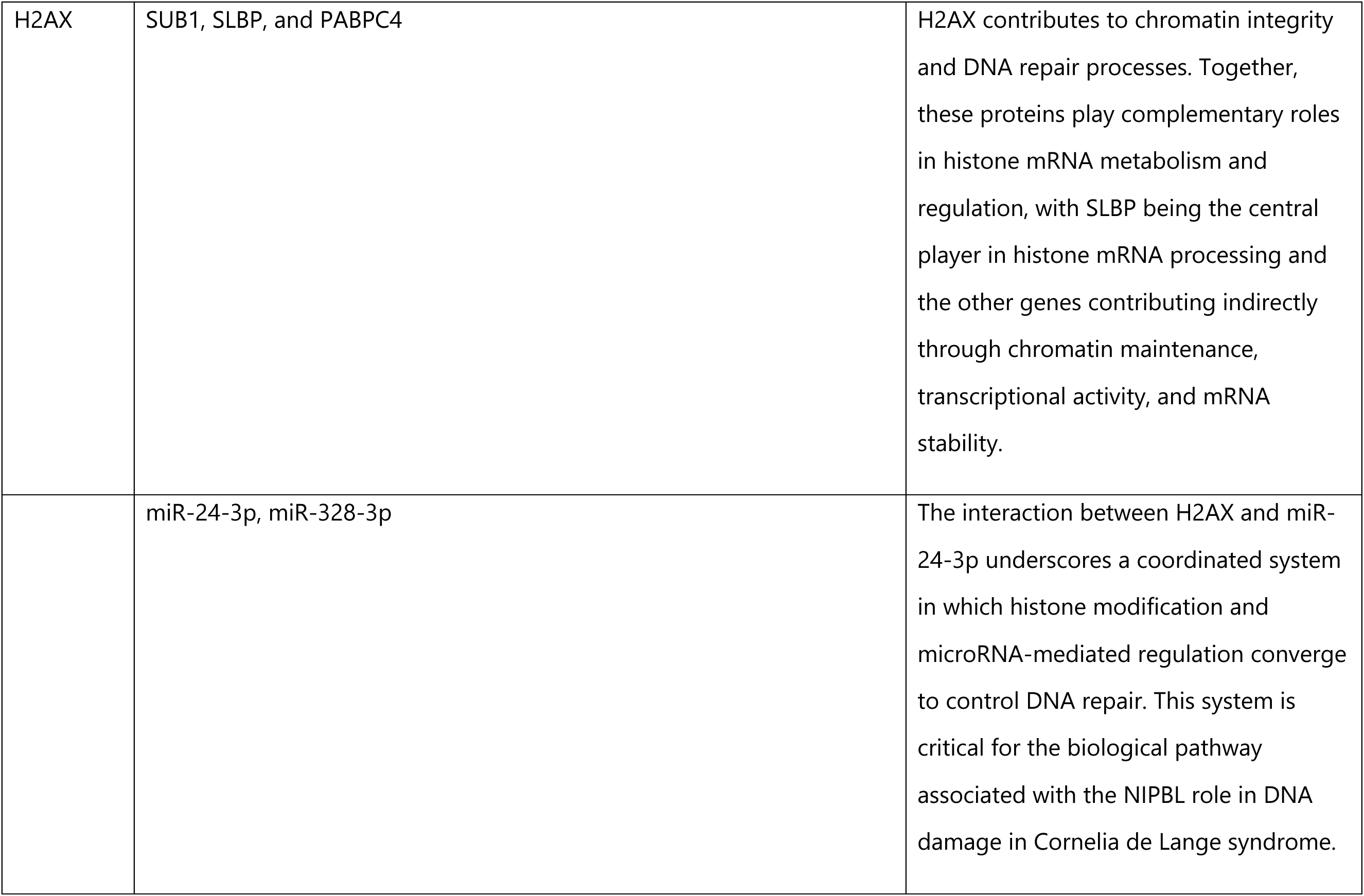
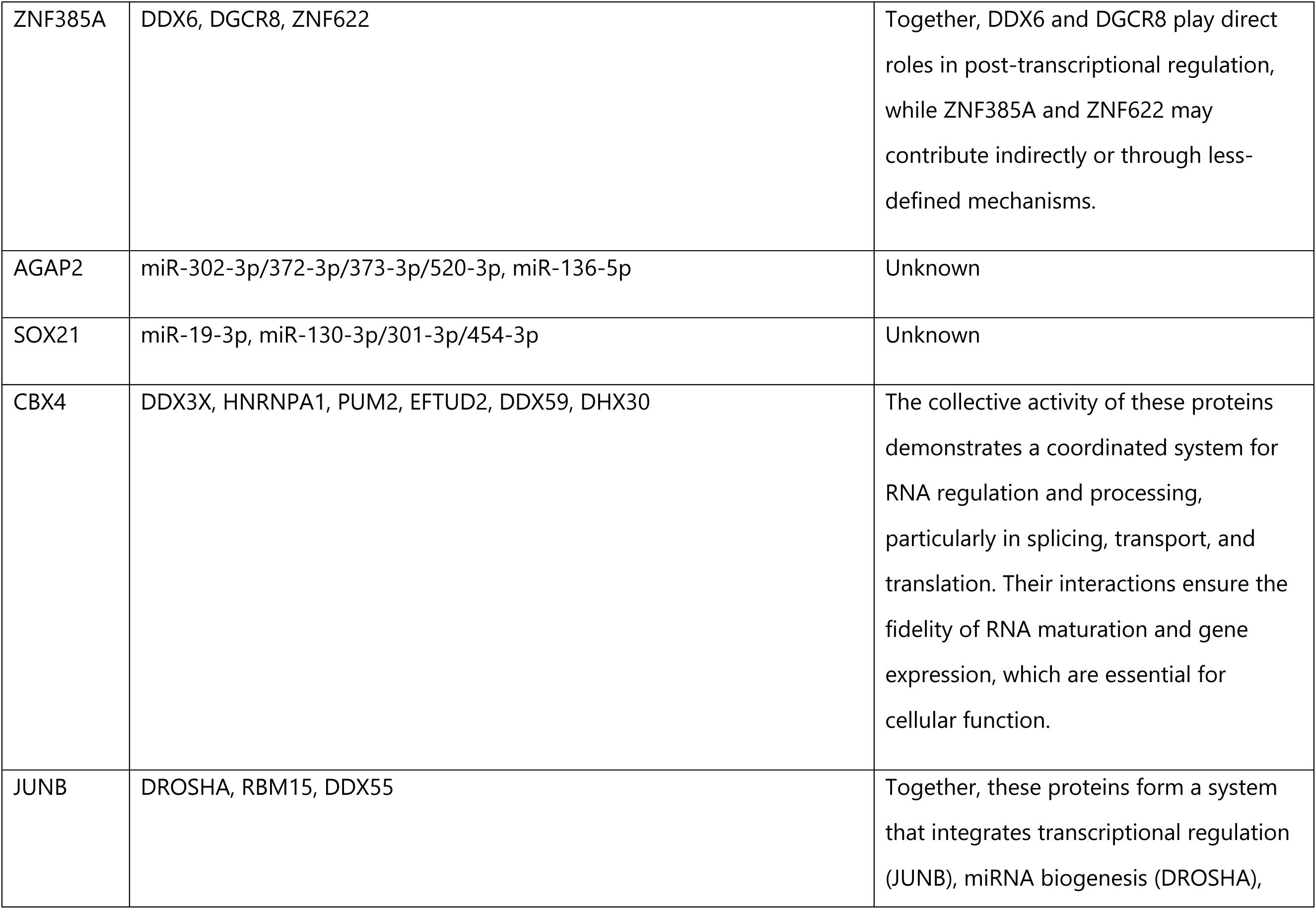
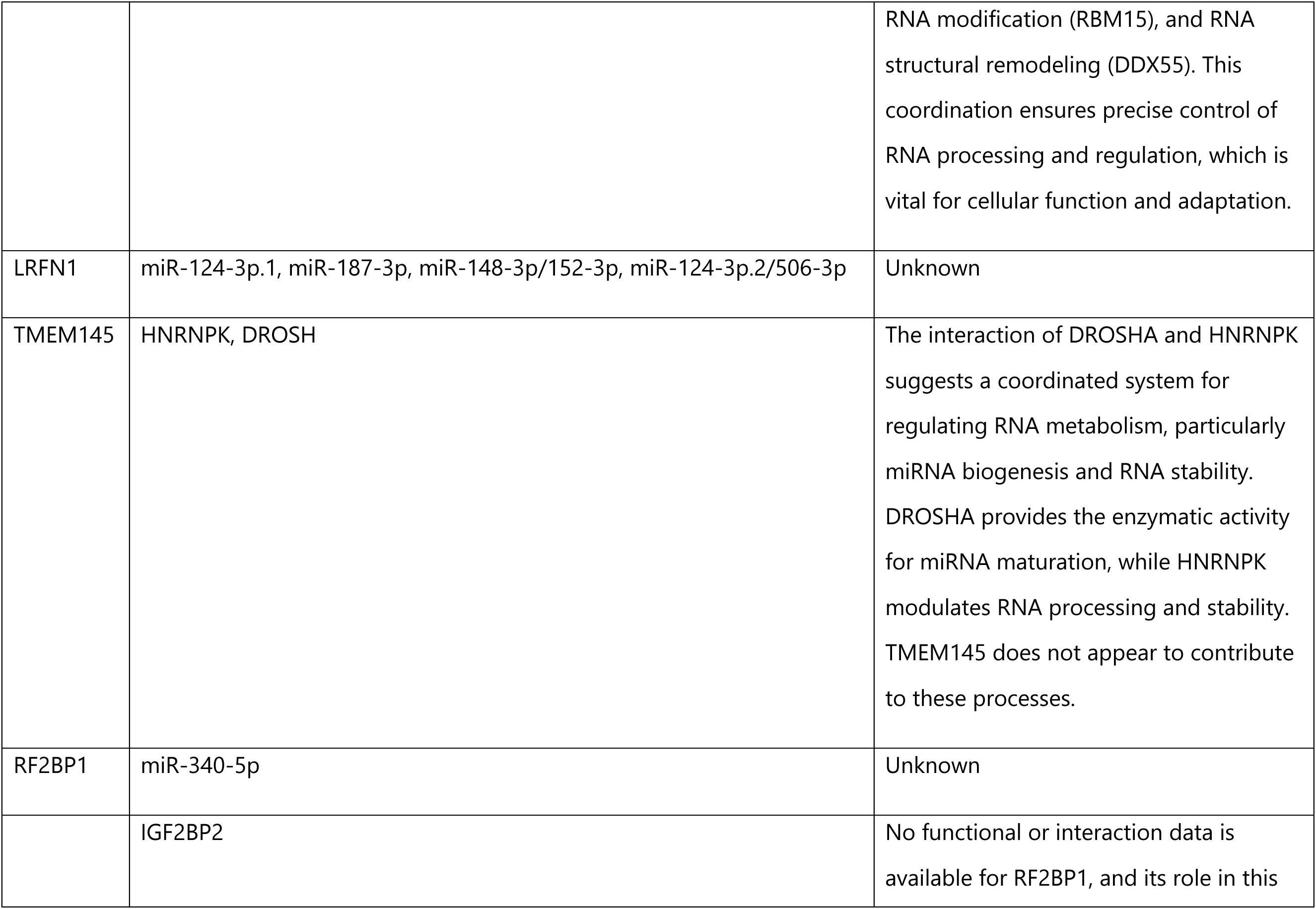
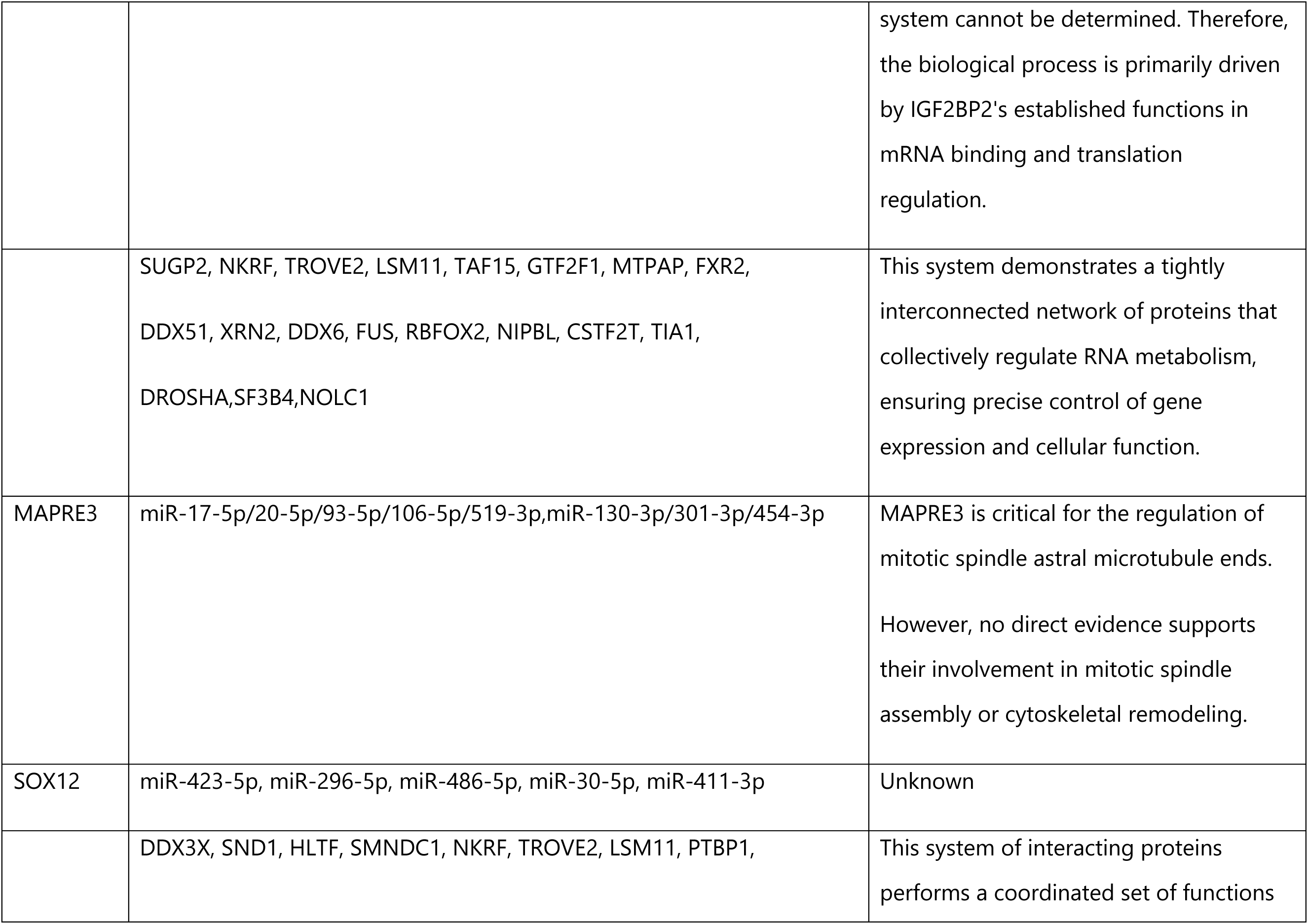
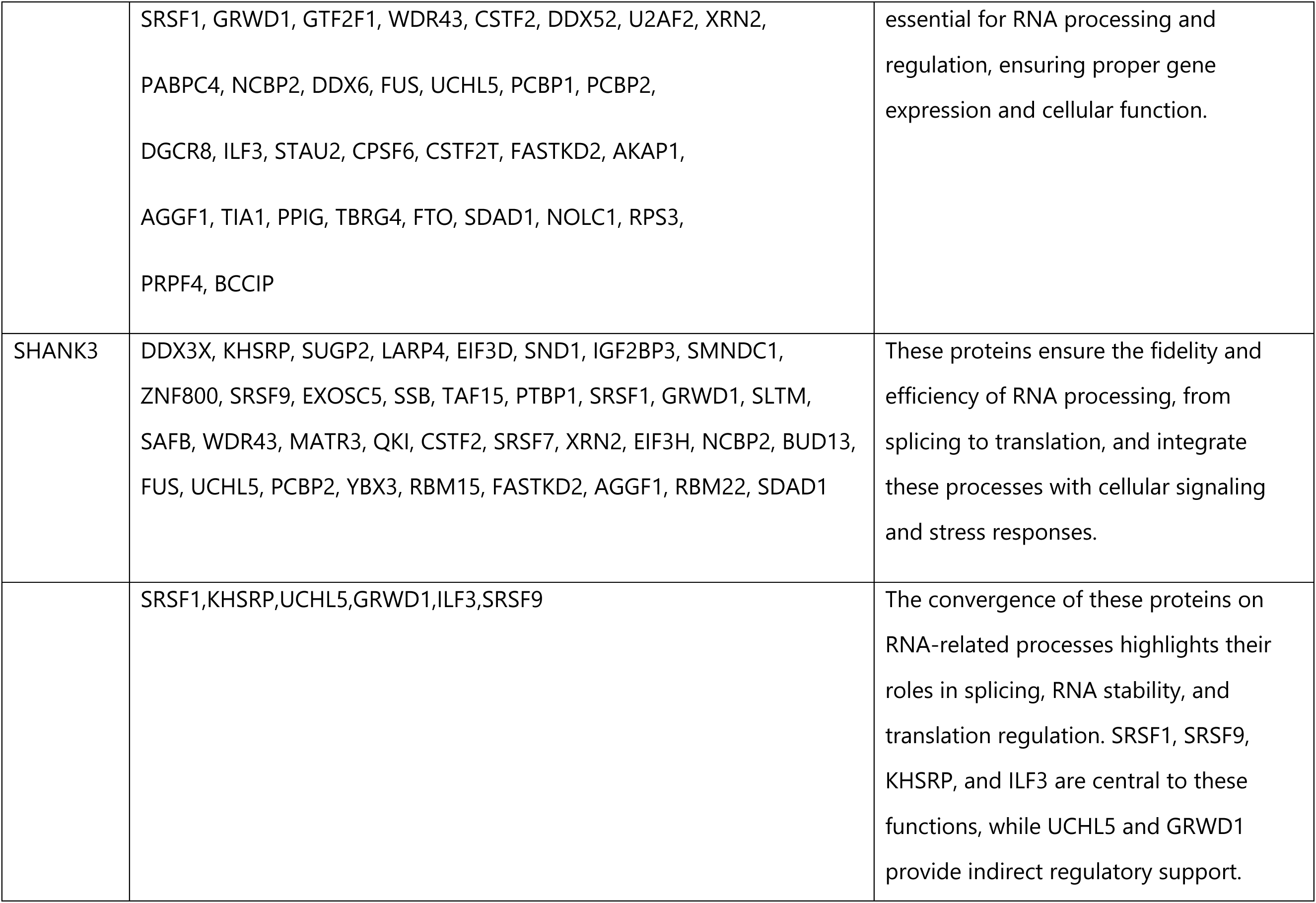
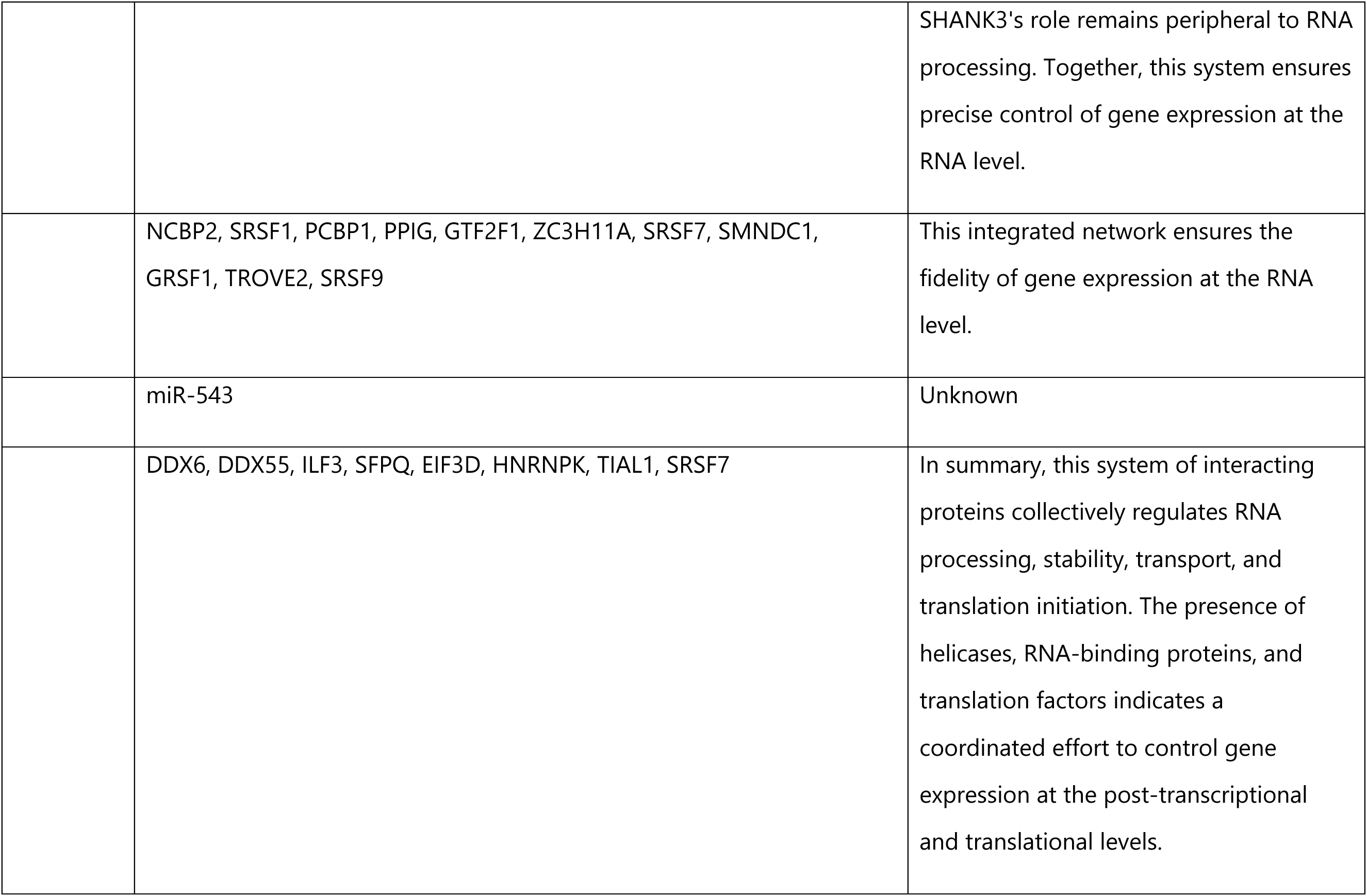
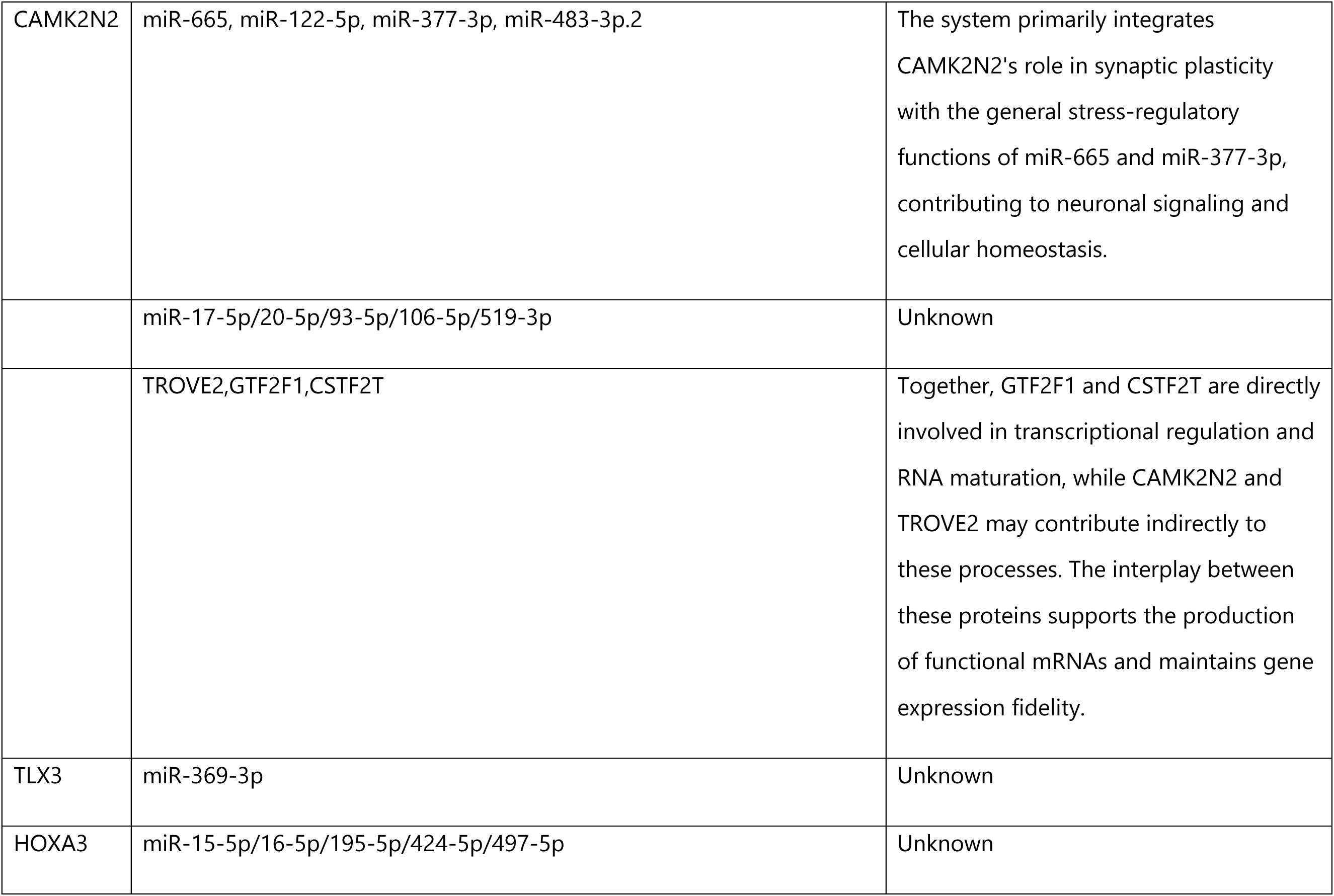
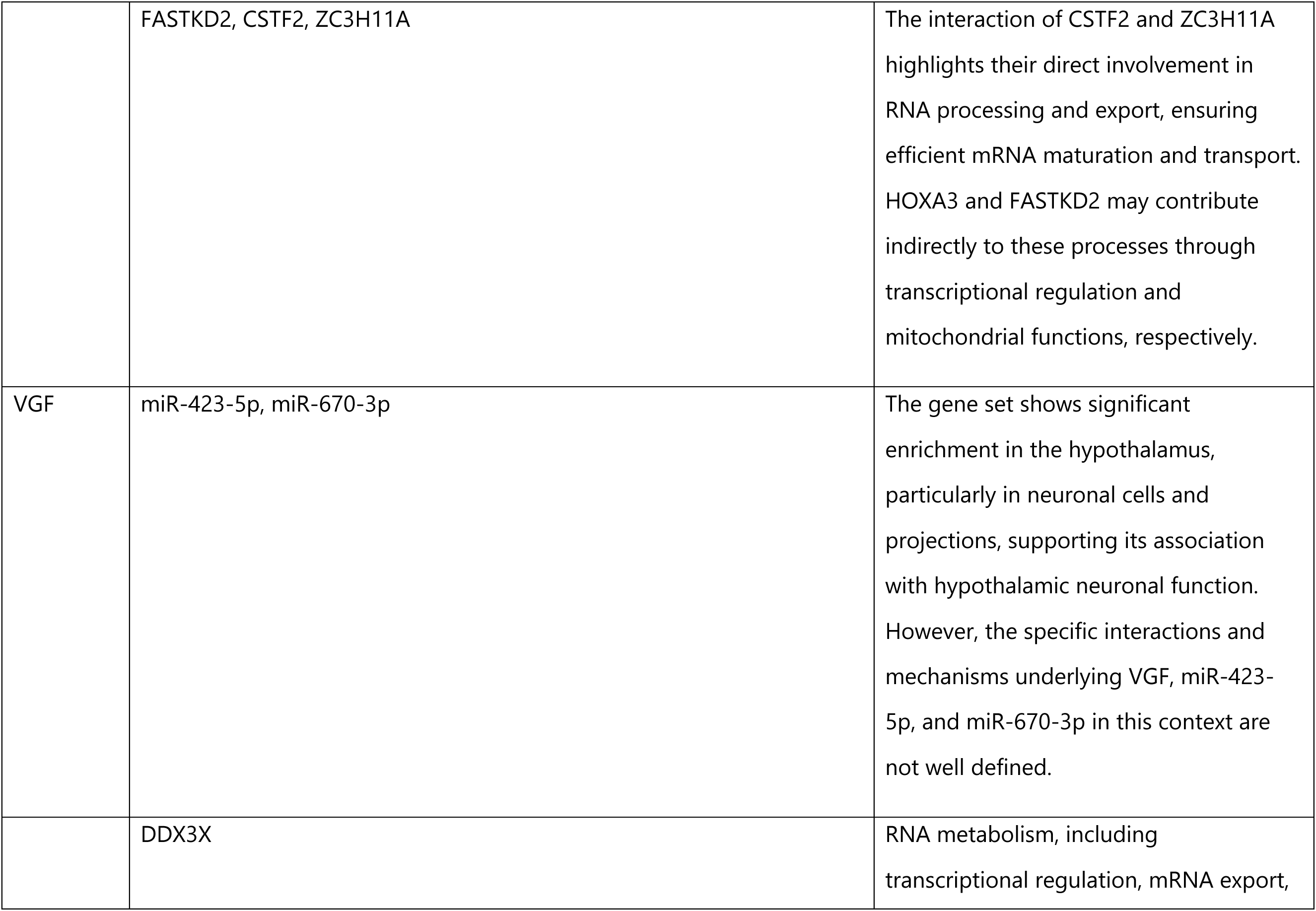
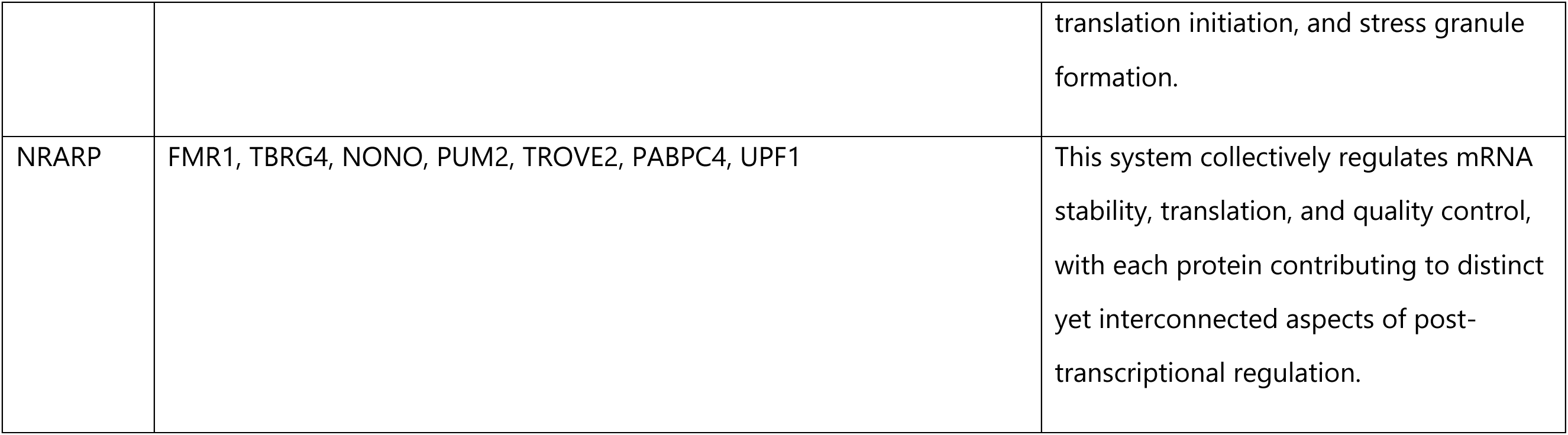

## DISCUSSION

In this study, we have identified approximately 3, 000 CFs (250 bp, ≥90% identity) in the 3’UTRs of ∼1, 500 genes across three to five diverse mammals. Out of which, 30% of them are at least 100 bp long. They are enriched with AT, TA, and TT with respect to other sequence types: 5’UTR, CDS, and 3’UTR. And they are not composed of low-complexity or simple repeating sequences. In fact, their complexity is similar to that of CDSs and 5’-/3 ’UTRs. As such, they are not the result of chance.

CFs are found with a higher concentration at the two ends of 3’UTRs. With scant evidence at this moment, we speculate that these CFs might contribute to preventing translation readthrough and polyadenylation, as AU-rich patterns resemble stop codons (UAA, UGA, and UAG), and canonical poly(A) signals (AAUAAA and AUUAAA).

Using a transformer-based genomic foundational model, GROVER, we have successfully identified distinctive tokens that distinguish CFs from the 3’UTR background. Based on the DNA embedding of the GROVER model, we have determined that signatory tokens are generally AT-rich, whereas non-CFs are G- and C-rich (Table 3). This information is essential for 3’UTR researchers, as AU-rich elements (AREs) in 3’UTRs are targets of AU-rich binding proteins, e.g., TTP and HuR (ELAVL1), which are most studied. A study has shown that translation efficiency can be elevated by introducing AREs (27, 55). The HuR family has been reviewed recently to be associated with neurological disorders (56). In addition to evolutionary conservation, we complement the functional annotation quest by examining variation intolerance in humans. Using two stringent, independent metrics, 36 CFs from 25 genes were identified as under intense negative selection pressure. These genes are strongly associated with the central nervous system, involved in neurodevelopment, and regulate RNA transcription. Based on eCLIP RNA-binding data and miRNA targets, RBPs and miRNAs that bind to the intolerant CFs play roles in transcription regulation and RNA metabolism. As RBP and miRNA targets are usually short (<10 bp), the length of CFs may suggest combinatorial interactions by these factors, enabling diverse regulatory mechanisms (19).

RBP and miRNA bindings are influenced by RNA secondary structure. We have explored the predicted secondary structure of CFs by mapping CFs to folded full-length mRNAs produced by RNAfold (57). And then the folding structures were fed into a convolutional neural network-based binary classifier to identify distinctive folding. However, no consistent features were found (data not shown). Further investigation is needed.

In conclusion, this study suggests that CFs play essential neurodevelopmental roles in 3’UTRs. Previous studies of UCEs primarily focus on their enhancer function. A recent computational study hypothesizes a role for them in homologous DNA pairing (58). Our work establishes a link between CFs and RNA processing, implicating a critical role in neurodevelopment mediated through RBPs and miRNAs. These insights pave the way for high-throughput functional screens, similar to established enhancer assays (23), to advance our understanding of the biological functions of ultraconserved elements.

## ACKNOWLEDGEMENTS

We acknowledge internal funding from Lafayette College.

## AUTHOR CONTRIBUTIONS

Eric S. Ho: Conceptualization, Formal analysis, Methodology, Validation, Writing—original draft, review, and editing. Nathan Dinh: Transformer-based binary classifier. Ash Baeck-Hubloux and Ciarra Troy: Enrichment analysis. Ava Severino: gene analysis and homology.

## SUPPLEMENTARY DATA

Supplementary Data are available at NAR online.

## FUNDING

This research received no external funding.

DATA AVAILABILITY

